# Haptenic adducts of β-lactam antibiotics elicit antibody responses with narrow clonality and specificity

**DOI:** 10.1101/2023.11.02.565155

**Authors:** Lachlan P. Deimel, Lucile Moynié, Guoxuan Sun, Viliyana Lewis, Abigail Turner, Charles J. Buchanan, Sean A. Burnap, Carolin M. Kobras, Mathew Stracy, Weston B. Struwe, Andrew J. Baldwin, James Naismith, Benjamin G. Davis, Quentin J. Sattentau

## Abstract

Many classes of small-molecule drugs form protein adducts *in vivo*, which may elicit antibodies via a classical hapten-carrier-type response, with implications for both allergy and drug sequestration. Although β-lactam antibiotics are a drug class long associated with these phenomena, the molecular determinants of drug-protein conjugation and consequent drug-specific immune responses remain incomplete. Here, we interrogated factors influencing penicilloyl adduct formation and immunogenicity, and used penicillin G (PenG) to probe the B and T cell determinants of drug-specific IgG responses in mice. We identify through deep clonotyping a dominant murine penicilloyl-specific clonal antibody class encompassing phylogenetically related *IGHV1*, *IGHV5* and *IGHV10* subgroup gene segments. Through protein NMR and x-ray structural analysis, we determined that adduct specific antibody clones—the MIL series—predominantly recognise the variable side-chain moiety (which for PenG is phenylacetamide) via a hydrophobic pocket, while secondary H-bond contacts with both thiazolidine and the adducted lysine residue is made. As a result, the cross-reactivity against other β-lactam antibiotics is limited. These data demonstrate the relationship between the chemistry of protein-reactive drugs such as penicilloyls, and how their predisposition to generating B cell responses can inform the functional implications at the clonal level.

**Highlights:**

- PenG readily forms immunogenic adducts on lysine sidechains of diverse self- and non-self proteins including complete serum under physiological conditions.
- PenG-protein adduction *in vitro* or *in vivo* is sufficient to elicit penicillin-specific IgG responses.
- Murine B cell clonotypic responses are characterised by near-uniform antibody binding modes of similar immunogenetic origin.
- The dominant murine PenG-specific clonotype is dominated by benzene ring recognition and correlates with serological cross-reactivity profiles.

## Introduction

Non-protein, low molecular weight compounds are typically non-immunogenic to the mammalian immune system. As exemplified by classical hapten-carrier biology, antibody responses against small molecules such as 4-hydroxy-3-nitrophenol acetyl (NP) require conjugation to a suitable carrier protein (*1*). The attachment of such a hapten to protein means that, within the same antigenic complex, cognate B cell receptors (BCRs) may cross-link a protein containing at an epitope encompassing the hapten, while associated peptidic components are presented to T helper (Th) cells, imparting linked help to propagate an activated hapten-specific B cell population (*2*– *4*).

In principle, these mechanisms may extend to small-molecule drugs, specifically those with reactive functional groups that may drive covalent conjugation with endogenous proteins under physiological conditions (*5*). One reported example of this phenomenon is for β-lactam antibiotics, such as penicillin G (PenG) (*6*). The β-lactam group of PenG is electrophilic and is the source of its inhibitory activity. However, it may also drive background reactivity with other biological nucleophiles leading to reactivity with primary amine-containing sidechains of lysine and potentially other nucleophilic residues including arginine, histidine, and cysteine under specific buffer conditions (*7*, *8*). Protein-PenG complexes are the antigenic determinants of antibiotic hypersensitivity.

The mechanistic underpinnings of the hypersensitivity reaction are immunologically heterologous, with the most common and well-characterised being T helper (Th) cell-mediated (type IV) that may be elicited in up to 30% of the population (*9*–*13*). However, the most clinically severe forms of drug hypersensitivity are antibody-mediated, particularly IgE-induced anaphylaxis (*14*). IgG-mediated hypersensitivity is less severe but relatively common (*14*, *15*). Penicillin is one of the most frequent causes of anaphylaxis and anaphylaxis-related deaths in humans (*15*). However, penicillin allergy diagnosis is currently highly inaccurate. Nearly 6% of the general population in the UK are allergic to penicillin according to their medical records, yet more than 95% of patients labelled as having a penicillin allergy ultimately can tolerate this class of drug (*16*). Patients with a penicillin allergy record have an increased risk of *Clostridioides difficile* and Methicillin-resistant *Staphylococcus aureus* infections and death; this is presumably through increased use of alternatives to β-lactam antibiotics (*17*). Furthermore, penicillin allergy diagnosis is associated with higher numbers of total antibiotic prescriptions (*18*), undermining antimicrobial stewardship goals and increasing the risk for antimicrobial resistance (*19*). A better understanding of the basis of penicillin hypersensitivity is needed to help predict which antibiotic recipients are, or will become, allergic (*20*, *21*).

Notably, although the first descriptions of penicilloyl-directed serological responses were made in 1961 (*8*), the relationships between the following phenomena remain incompletely understood: i) the biochemical basis of PenG–protein adduction *in vivo* and *in vitro*; ii) the relative immunogenicity of fully chemically-characterised and purified PenG adducts; iii) the immunophylogenetics of B cells specific to PenG- protein complexes; iv) the structure/function characteristics of antibody clones specific to these adducts. These findings may serve as a template for understanding allergic outcomes in the clinic, suggest approaches for rational prediction of hypersensitivity and potentially inform the design of antibiotics that avoid sensitisation.

Here, as part of such a roadmap to understand the antigenic and immunogenic determinants of PenG-specific antibody responses, we describe the systematic investigation of the relationship between the protein-conjugating chemical properties of PenG and its immunogenicity in a mouse model. We carried out clonal B cell analyses and performed complementary structural and biochemical characterisation of the clonotypic PenG-specific antibody response in mice. These findings offer a rational basis for understanding anti-drug antibody (ADA) responses, revealing how drug conjugation chemistries can affect the downstream antibody response.

## Results

### Immunogenicity of pre-complexed penicillin-protein antigen

PenG has constituent β-lactam, thiazolidine and benzylamide sidechain moieties (**Fig 1a**); the β-lactam ring is long known to react with nucleophilic protein sidechains, particularly the ε-amino groups of lysine residues, (*3*, *7*, *8*, *22*–*24*) (**Fig 1b**). To probe PenG-protein conjugation efficiency, we titrated adduct formation on the model protein hen egg lysozyme (HEL). HEL was chosen for being a relatively small and stable protein (∼14 kDa, 129 a.a.) with 6 lysine residues. Various buffer, drug amounts and pH conditions were tested *in vitro*, and global site-specific drug occupancy was evaluated via mass spectrometry (MS). These revealed pH- and buffer-modulated conjugation levels (**Fig S1a**; **Document S1**). Mapping of adduct formation through tryptic digest followed by site-specific liquid chromatography tandem MS (LC-MS/MS) analysis confirmed adduct formation at lysine residues 1, 13, 33, 96/97 and 116 (**Fig S1b**). The adduct-associated mass change is consistent with previous reports of β-lactam folding (**Fig S1c**). Based on these data, we then used 0.1 M Na2CO3 pH 8.0 (a close-to-physiological pH) for *ex vivo* conjugation of PenG to various recombinant carrier proteins for subsequent immunisation.

**Figure 1:**
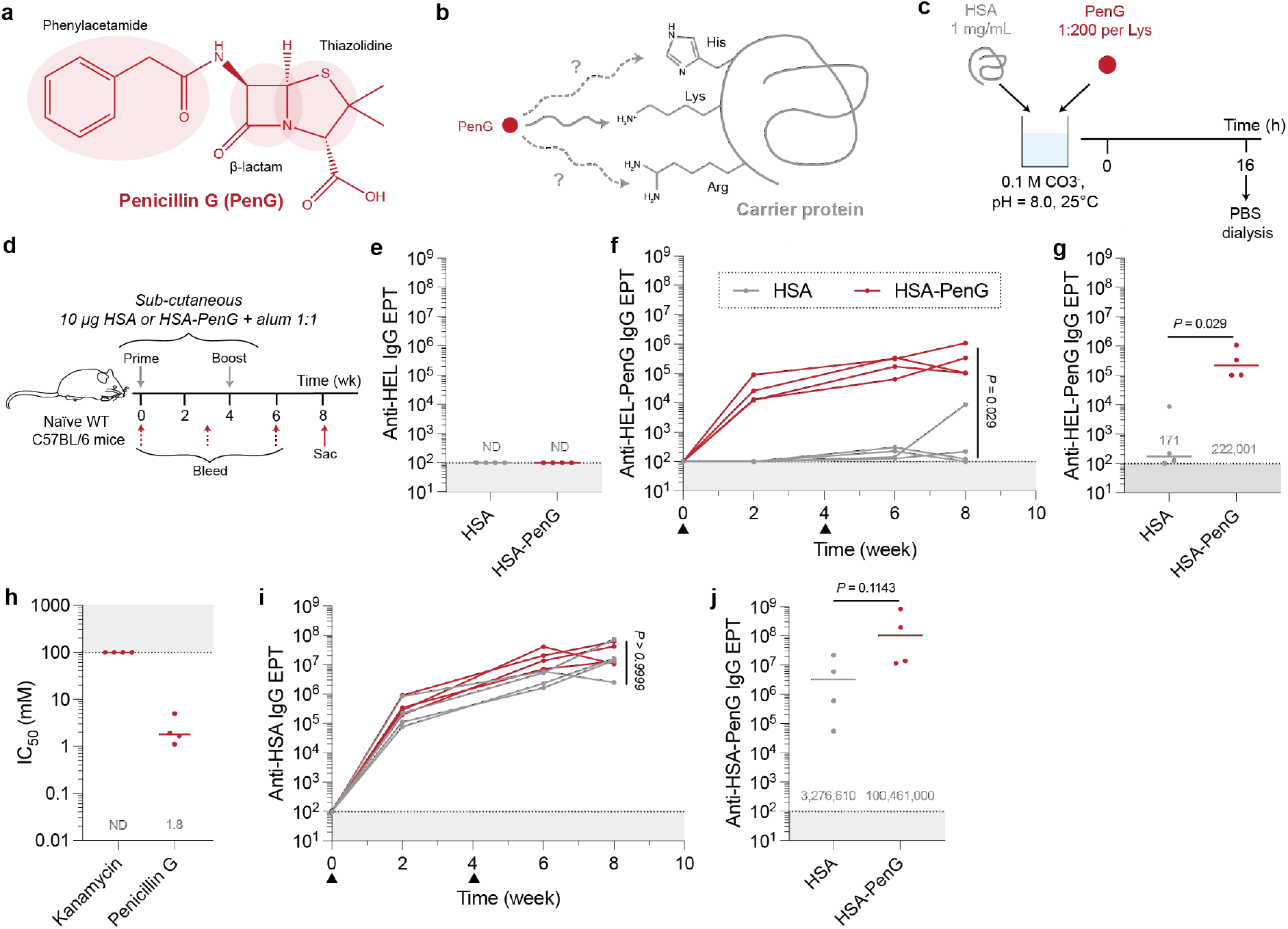
Chemical characterisation and serological responses against PenG-derived penicilloyl adducts. **(a)** Chemical structure of PenG. **(b)** Proposed target residues of electrophilic B-lactam; primary reported target of lysine with potential targeting of arginine and histidine. **(c)** 1 mg/mL HSA and PenG (1:200 per Lys) were mixed *in vitro* with 0.1 M HCO3-, pH = 8.0. This was left at 25°C for 16 h before dialysis into PBS. **(d)** Sex-matched 6-week-old naïve WT C57BL/6 mice were twice immunized (wk 0 and 4) with 10 µg HSA or HSA-PenG in alum. **(e)** Terminal HEL-specific IgG EPT was evaluated. PenG-specific endpoint titres were evaluated by screening cross-reactivity against HEL-PenG. IgG titres against PenG were evaluated both **(f)** longitudinally and **(g)** at the terminal timepoint. **(h)** Competition ELISA was conducted, wherein HSA-PenG antisera binding for HEL-PenG was competed out with soluble PenG. **(i)** Longitudinal protein backbone-specific, HSA, and **(j)** terminal HSA-PenG-specific IgG endpoint titres were screened. Dots represent data from a single animal. Groups were compared via Mann-Whitney test.

Pilot immunogenicity analysis of HEL-PenG formulated in aluminium hydroxide (alum) adjuvant was conducted by subcutaneous (s.c.) administration to C57BL/6 mice. Antisera were titrated by ELISA on an unrelated PenG-modified protein (human serum albumin-PenG; HSA-PenG), which revealed modest but significant (*P* < 0.05 Mann Whitney U) isotype-switched IgG responses raised against the penicilloyl adduct (**Fig S1d–g**). Since HEL is a weak Th cell antigen in mice (*25*, *26*), we next evaluated the antibody response generated by PenG pre-complexed to the more antigenic HSA using the conditions optimised for HEL (**Fig 1c**). Site-specific occupancy of penicilloyl adducts was evaluated via LC-MS/MS and diverse lysine occupancy was observed (**Fig S2**). Mice were immunised with HSA or HSA-PenG formulated in alum, followed by periodic blood sampling (**Fig 1d**). Anti-penicilloyl serum IgG responses were measured by ELISA against HEL-PenG; no IgG cross-reactivity was detected against unmodified HEL (**Fig 1e**). Post-prime, HSA-PenG antisera displayed considerable IgG reactivity with HEL-PenG, whereas no reactivity was detected in the HSA-alone antiserum (**Fig 1f**). Together, these suggest an anti-PenG-adduct-specific response. Post-boost and at the terminal timepoint, the median HEL-PenG-specific IgG endpoint titre (EPT) was ∼2.2 x 10^5^ for the HSA-PenG antisera and ∼1.7 x 10^2^ for the HSA antisera (*P* = 0.029, Mann-Whitney test) (**Fig 1g**).

These antibody responses were generated against a protein-conjugated PenG derivative. Characterisation was consistent with direct β-lactam opening, but we cannot discount a pathway involving intermediates of penicillanic acid (**Document S1**) (*23*, *24*). HSA-PenG antiserum binding to HEL-PenG was out-competed by free PenG, with a median IC50 of 1.8 mM, while kanamycin (an unrelated non-β-lactam-type antibiotic) failed to detectably compete for antibody binding (**Fig 1h**). These data reveal that antibodies raised against the autologous PenG adduct are cross-reactive with free PenG. Administration of HSA-PenG did not affect the antibody responses against the protein unmodified protein backbone, compared with mice immunised with HSA alone (*P* > 0.9999, Mann-Whitney test) (**Fig 1i,j**).

### Self-protein carriers elicit penicillin-specific antibodies

Having demonstrated strong B cell immunogenicity of PenG complexed with foreign protein (HEL or HSA) backbones, we next tested a native and otherwise tolerogenic self-antigen, mouse serum albumin (MSA), as a more physiologically-relevant model (*27*). First, pure MSA was pre-complexed with PenG in buffer under the optimised conditions, as described previously. Animals were immunised three times (wk 0, 4 and 8) with or without alum and terminal (wk 10) IgG antibody titres were evaluated by ELISA against OVA-PenG (**Fig 2a**). 5/6 (83%) animals immunised with MSA-PenG in alum elicited a detectable serum endpoint titre against the adduct (median titre of ∼ 5.2 x 10^3^) and 3/8 (38%) responders in those immunised with MSA-PenG without adjuvant (**Fig 2b**). These data show that extrinsic adjuvantation is not required to elicit anti-adduct IgG responses even when presented on a modified self-protein. By contrast, no animal immunised with unmodified MSA with or without alum gave a detectable PenG-specific response (*P* = 0.033, Kruskal-Wallis test) (**Fig 2b**). Although the MSA-PenG-elicited PenG-specific antibody titres were lower compared with HSA- PenG (**Fig 1a**; **Fig 2b**), these data show that a penicillin-specific antibody response can be generated even when using a highly T-immunorecessive backbone, and that self-proteins can act as immunogenic carriers. No reactivity was detected against OVA in any group (**Fig 2c**).

**Figure 2:**
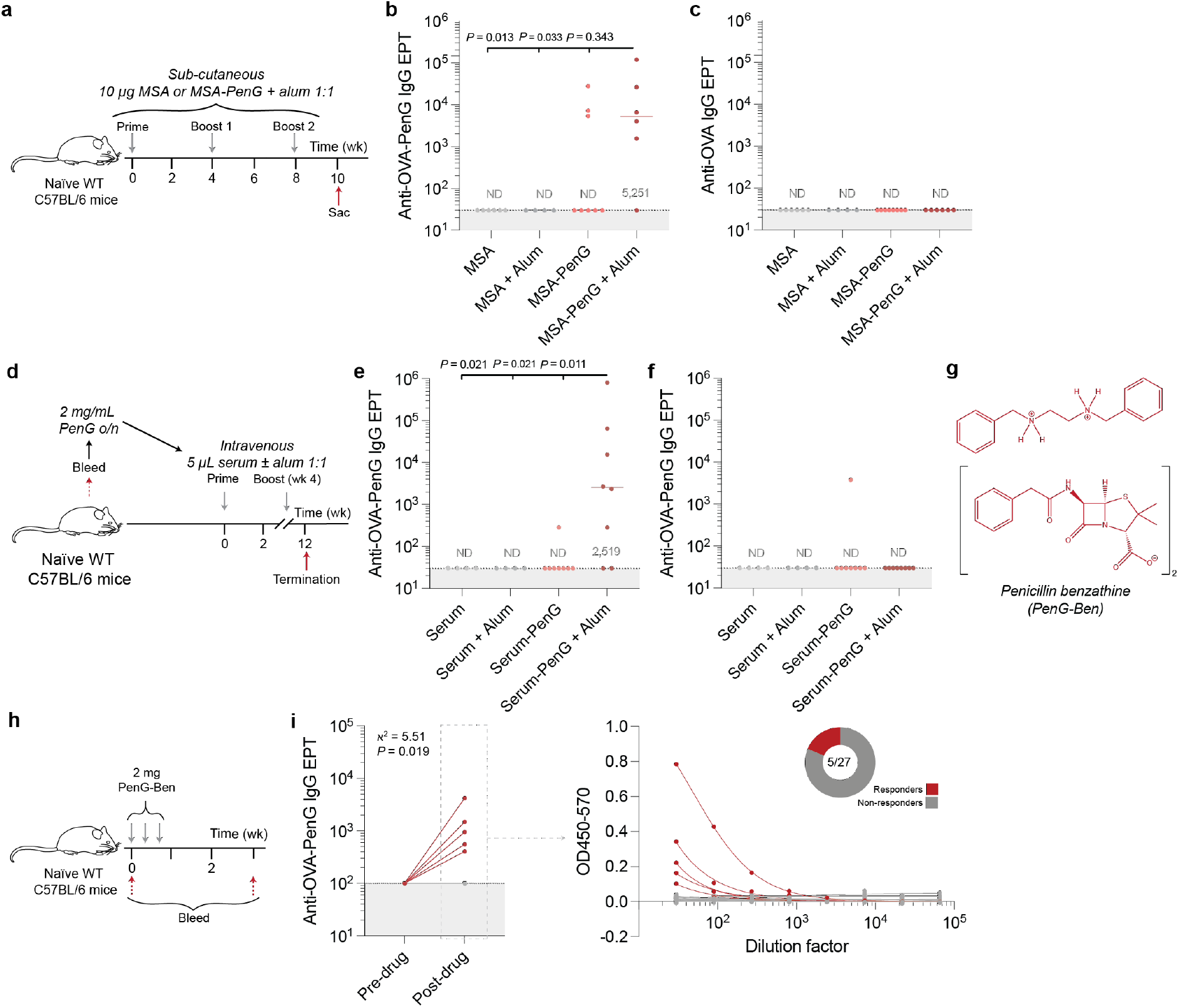
Factors affecting the PenG-specific serological response. **(a)** Sex-matched C57BL/6 mice were immunized three times and the serological response at the terminal timepoint (wk 10) was evaluated. **(b, c)** Mice were immunized three times (wk 0, 4 and 8) and the terminal (wk 10) IgG PenG-specific EPTs were evaluated. **(d)** Mice were bled and 2 mg/mL PenG was added and mixed end-to-end overnight. Animals were subsequently immunized (wk 0 and 4) intravenously with seum-PenG with or without alum. **(e,f)** Terminal IgG EPTs were evaluated. Data were compared via a Kruskal-Wallis test. **(g)** PenG-Ben structure. **(h)** Mice were given PenG-Ben intramuscularly. **(i)** PenG-specific IgG titres were evaluated. Data were compared to pre-administration, evaluating the ratio of responders via Chi-squared test.

Finally, to develop an *ex vivo* model for self-protein:PenG adduct formation and immunogenicity, we tested the ability of mouse serum itself to act as a carrier for PenG. PenG-naïve mice were bled, serum isolated and incubated *ex vivo* with 2 mg/mL PenG for 16 h. The resulting serum-adduct mixture was then administered intravenously (i.v.), into the autologous mice, with and without alum, to mimic a clinical route of penicillin administration (**Fig 2d**). Terminal IgG endpoint titres were evaluated against OVA-PenG, which revealed 6/8 (75%) of animals primed and boosted with adjuvanted serum-PenG responded with a detectable IgG titre against OVA-PenG and a median titre of ∼ 2.5 x 10^3^ (**Fig. 2e,f**). Interestingly, only a single animal that received unadjuvanted serum-PenG gave a detectable PenG-specific IgG titre, implying that adjuvantation, most likely provided by bacterial infection during therapeutic penicillin use, is probably required under these conditions to overcome a threshold for immunogenicity.

### Immunogenicity of free penicillin

Having shown that PenG is immunogenic when pre-complexed with diverse protein carriers including mouse serum, we next evaluated whether free penicillin, as would be administered in the clinic, might be sufficient to induce an antigen-specific antibody response. First, we tested whether PenG delivered i.v. daily to mice in 2 x 1 week-long courses was immunogenic (**Fig S3a**). However, no IgG or IgM drug-specific responses were detected when compared to mice given control PBS (**Fig S3b,c**). Second, the immunogenicity of orally-administered antibiotic was evaluated, using the oxo-homologue penicillin V (PenV). Unlike PenG, PenV is used for oral administration as it does not degrade under the acidic conditions of the stomach (*28*). Mice were given PenV *ad libitum* for two 3.5-day intervals. Some mice were additionally given an i.p. dose of lipopolysaccharide (LPS) (0.5 mg/kg) to mimic the systemic increase in endotoxin expected from a bacterial infection, where antibiotics such as PenG/V would be clinically used (**Fig S3d**). Despite this, no specific PenG titres were observed (**Fig S3e**). Notably, administration of LPS increased the background reactivity of serum: mice given PenG and LPS or drug-free water and LPS-only both exhibited modest reactivity against HSA-PenG, which we attribute to induction of B cells producing polyspecific IgG responses (*29*).

We further hypothesised that the failure to initiate a serological response following the administration of free drug may be a consequence of rapid clearance of the antibiotic in a mouse; for instance, mice have an extremely high cardiac output (*30*) and renal clearance of PenG is efficient (*31*), with a previously reported half-life of approximately 15 mins. Fast clearance will restrict endogenous PenG adduct formation in this animal model, ultimately reducing the probability for antigen-B cell encounter and BCR cross-linking. Therefore, to extend the availability of free drug *in vivo*, penicillin G benzathine (PenG-Ben) was used. This is a formulation of PenG with diphenylethylene diamine that renders PenG poorly soluble for use in slow-release delivery (**Fig 2g**). When administered intramuscularly, PenG-Ben is solubilised over days to weeks, to release PenG systemically (*32*). PenG-Ben was administered intramuscularly (i.m.) to 27 mice (**Fig 2h**). After the antibiotic course, 5/27 (18%) mice generated IgG responses to OVA-PenG, which is a significant proportion when compared with the pre-immune serum (χ2 = 5.51, *P* = 0.019) (**Fig 2i**). Reactivity against unmodified OVA was not detected (**Fig S3f**), suggesting a specific but infrequent PenG-Ben-specific antibody response following PenG-Ben administration in mice.

### Cross-reactivity of anti-penicillin IgG responses

Since penicillin shares some structural homology with other β-lactam antibiotics, we screened the cross-reactivity of the HSA-PenG serological response. Ovalbumin (OVA) was modified with a set of penicillin antibiotics with differing side chains, and with β-lactam antibiotics from other classes (including cephalosporins and carbapenems), using the previously determined conditions (**Fig 1c**). Autologous reactivity against OVA-PenG was the greatest of the diverse OVA-X panel tested, with a median IgG EPT of ∼ 2.5 x 10^6^ (**Fig 3a**). Interestingly, there was limited reactivity against OVA-ampicillin (median EPT of ∼ 2.4 x 10^3^), which differs only in its benzylic amine substituent, and similarly carbenicillin (median EPT of ∼ 4.9 x 10^3^), which differs by its benzylic carboxyl constituent. However, there was considerable cross-reactivity against OVA-oxacillin (median EPT of ∼ 8.5 x 10^4^), despite the greater variation in sidechain compared with ampicillin. These data suggest that the polyclonal response tolerates some change in side-chain but that this may also be blocked by simple alterations at pivotal sites (such as the ampicillin H→NH2, or the carbenicillin H→COOH change). A subset of 1^st^–4^th^ generation cephalosporin- and carbapenem-type antibiotics were also screened for cross-reactivity. HSA-PenG antisera displayed limited (albeit above the detection limit) cross-reactivity against these modified OVA antigens (IgG EPTs ∼ 10^3^). Since cephalosporins and carbapenems have differing β- lactam-encompassing cores (*33*), these data suggest that the PenG-raised antibody response is largely dependent on the 6-aminopenicillanic acid-derived core. Finally, since β-lactam antibiotics in part mimic the proteoglycan subunits D-Ala-D-Ala, it was of interest to assess reactivity with these repeating units via a competition ELISA (**Fig 3b–d**). However, competition for binding with soluble D-Ala-D-Ala was not detected.

**Figure 3:**
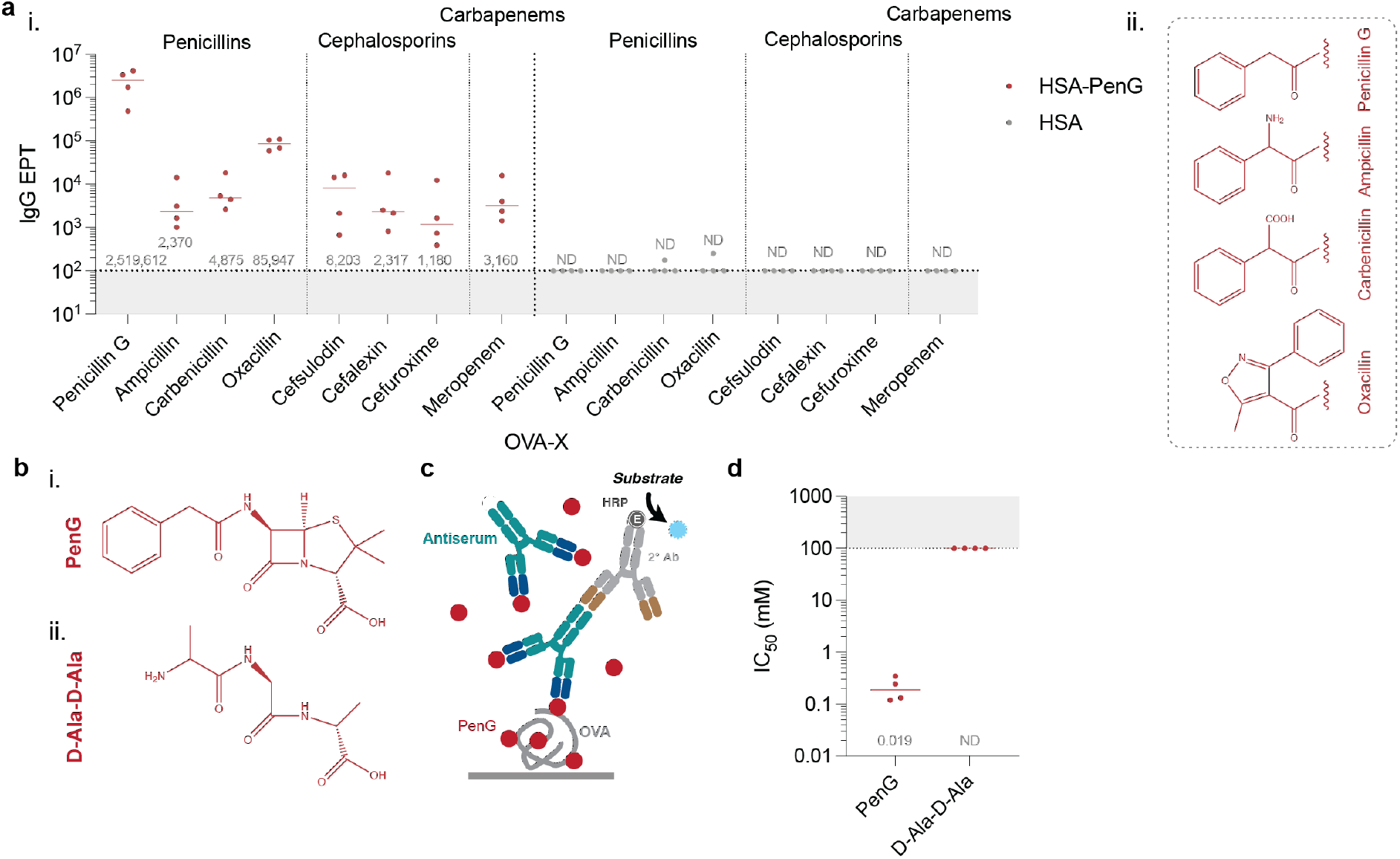
Specificity and cross-reactivity in the PenG-specific antibody response. **(a) i.** HSA-PenG antisera were screened against a set of β-lactam antibiotic-modified OVA. Data reflect the IgG EPT against the drug adducts. **ii.** Sidechains of penicillins tested. **(b)** Structural comparison between **i.** PenG and **ii.** D-Ala-D-Ala. **(c)** Competition ELISA format. **(d)** Evaluation of the relative competitiveness of D-Ala-D-Ala.

### Clonotypic B cell responses to PenG adducts

To evaluate the B cell response at the clonal and molecular levels, PenG-specific B cells were isolated and variable regions cloned using techniques previously described (*34*, *35*). Mice were immunised with HSA-PenG in alum and draining inguinal lymph nodes (iLN) were harvested 2 weeks post-prime (**Fig 4a**). To isolate PenG-specific B cells, molecular probes were synthesised by chemically modifying another carrier protein (HIV-1 gp120) that we have validated as giving low background and high specificity in a different hapten-carrier context (*25*). Gp120 was modified with PenG and then modified with fluorophore Alexa Fluor 647 using a corresponding NHS ester. We further tetramerised biotinylated gp120 with streptavidin-phycoerythrin. These antigen probes were used to sort the PenG-specific B cells on pre-gated non-naïve (DUMP^-^B220^+^IgD^-^) B cells (**Fig 4b**; **Fig S4a**). B cells were sorted from four mice and cell clonality was inferred according to the VH sequences (**Fig 4c**).

**Figure 4:**
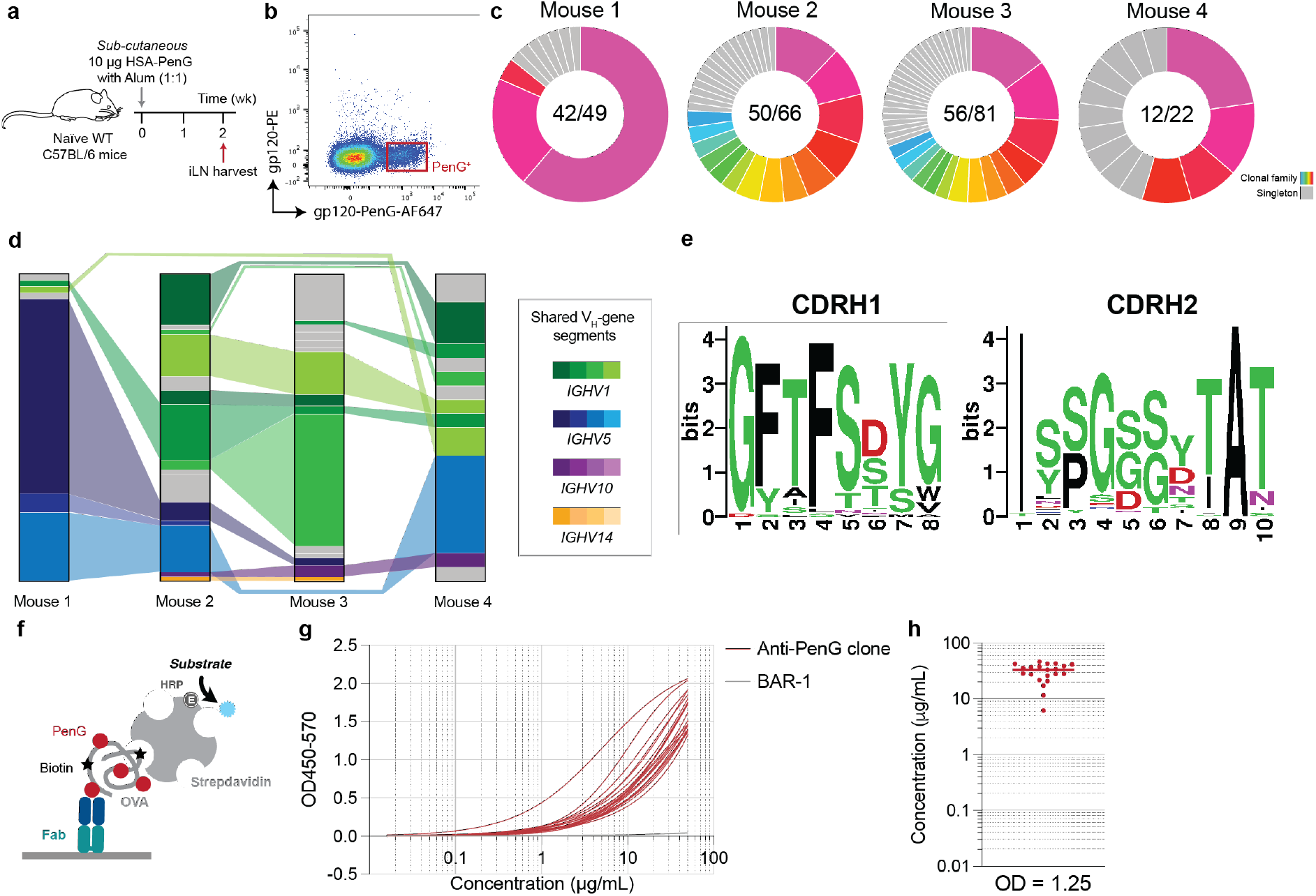
Immunogenetic characterisation of the PenG-specific response. **(a)** Immunisation schedule. **(b)** Antigen probe sorting strategy on pre-gated non-naïve B cells (DUMP^-^B220+IgD^-^). **(c)** Inferred clonal families from of the PenG probe-sorted B cells (coloured) and singletons (grey). **(d)** Inferred V_H_ gene segment across the sequenced B cells. Bar segment sizes proportional to the number of B cells of the same V_H_ origin. Joining of the segments connote the shared utilisation between mice. Segment colours reflect the *IGHV* subgroup. **(e)** Logo plots of the CDRH1 and CDRH2 amino acid sequences from all sequenced B cells in all animals. **(f)** Fab binding specificity assay setup. **(g)** ELISA trace for a subset of 28 Fabs and **(h)** their OD1.25 intersection (estimate for EC_50_) values. BAR-1 (grey) is an unrelated negative control specific to sialyllactose.

Interestingly, considerable sharing of VH gene segments was observed between the mice, suggesting that similar clonotypes were raised across animals (**Fig 4d**; **Fig S4b**). Notably, the VH gene segments utilised were from four highly phylogenetically-related subgroups: *IGHV2*, *IGHV5*, *IGHV10* and *IGHV14*. This striking homology is reflected in conservation of the CDRH1 and CDRH2 amino acid sequences across the mice (**Fig 4e**). Together, these data suggest that there are preferred structural and functional motifs encoded in these germline segments that facilitate binding with the drug. Immunogenetic analysis revealed a defined and ordered CDRH3; the length was bimodal, either short (5–6 aa) or long (17 aa) (**Fig S5a**). Short CDRH3s were dominated by an ARG motif for the first three residues, with a diverse C-terminal end, while the long class possesses a negative N-terminus, a neutral centre and a positive C-terminus (**Fig S5b**).

Clonal families were evaluated. The two largest families were isolated from mouse 1 and their germinal centre trees are shown (**Fig S6**). The largest clonal family (**Fig S6a**) exhibits a ‘clonal burst’ following the acquisition of the S11T mutation, a phenomenon reportedly associated with the acquisition of an affinity-improving mutation that renders the clone more competitive for antigen uptake and T cell help (*36*). This mutation was also found in a separate clade on the same tree. The T85S mutation was also identified twice on separate clades.

Finally, to validate the PenG specificity of the antibody response, a subset of V-region pairs from all mice and with diverse gene segment origins were cloned, and corresponding Fabs were expressed and purified. Binding was validated via ELISA; all Fabs bound the PenG adduct probe, while an unrelated antibody Fab (BAR-1) did not detectably bind (**Fig 4f–h**). These data confirmed that the sorting approach was highly specific.

### Structural, biochemical and biophysical characterisation of the antibody response to PenG

We selected a subset of PenG-specific clones with divergent CDRH3 lengths to further dissect the binding characteristics of the clonotypical response to penicilloyl adducts: MIL-1 (*IGHV5-6*01*; 17 a.a. CDRH3), MIL-2 (*IGHV5-17*01*; 6 a.a. CDRH3) and MIL-3 (*IGHV10-3*01*; 9 a.a. CDRH3). We designed a soluble Lys-PenG ligand as a reductionist adduct mimic reflecting β-lactam unfolding on the ε-amine of lysine residues (**Fig 5a**). The ligand was characterised by both high-resolution MS and NMR, which demonstrated a highly pure product (**Document S1**).

**Figure 5:**
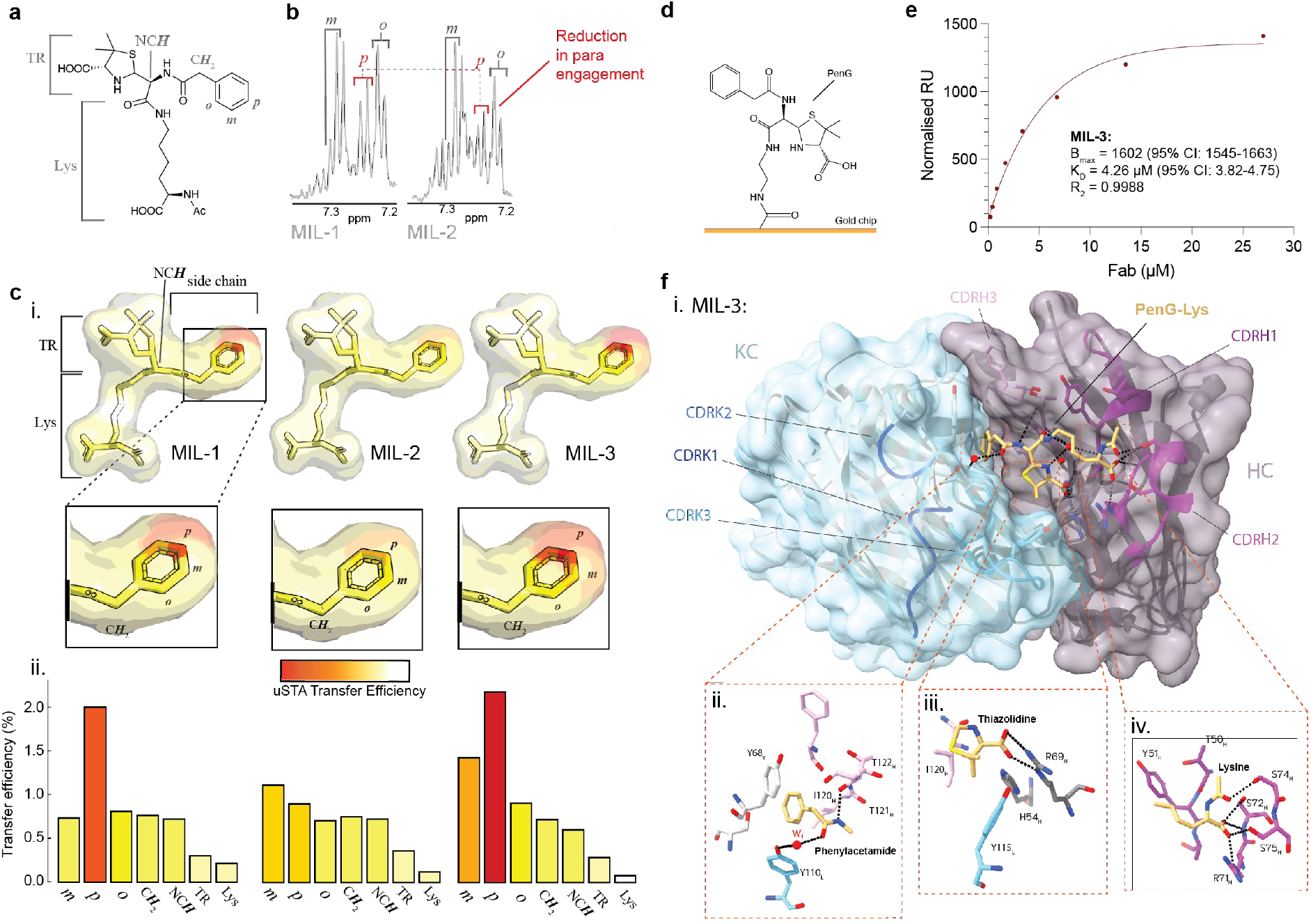
Structural, biochemical and biophysical characterisation of the PenG-specific response. **(a)** Annotated chemical structure of PenG-lys construct used for uSTA analysis. **(b)** Saturation transfer difference spectrum was generated from the difference between the raw off-resonance (gaussian at 37 ppm) and the raw on-resonance (gaussian at 9 ppm) spectra. Data showing differences in para engagement in the benzene ring of MIL-2 fab versus MIL-1. **(c)** i. Heatmaps corresponding to saturation transfer efficiency of PenG-lys (1 mM) with MIL-1–3 (5 µM). ii. Histographic saturation transfer efficiencies. Red indicates high transfer efficiency. **(d)** SPR chip design. **(e)** Biophysical characterization of MIL-3 via SPR. **(f)** i. Top view of the x-ray structure of the MIL-3 Fab bound to the PenG-Lys (beige sticks). Both heavy (monochrome plum) and light chain (monochrome blue) CDRs are marked. Key residues within 4.0 Å of ligand aspects ii. phylacetamide, iii. thiazolidine and iv. lysine are shown. Hydrogen bonds are shown as black broken lines. Water is marked in red.

First, atomic resolution of the binding pose was dissected using universal saturation transfer analysis (uSTA) (*25*, *37*). High transfer efficiencies were observed on the benzene ring of Lys-PenG with all three fab fragments, with the *p*-proton showing the most considerable engagement for MIL-1 and MIL-3 (**Fig 5b,c**). MIL-2 exhibited a lower transfer efficiency for the para-benzene proton compared with MIL-1 and MIL-3, however the binding was still mediated through the benzene ring. The lysine residue of the construct exhibited minimal transfer efficiency in all cases, implying, as indicated in earlier data, that binding is dominated by adduct rather than sidechain binding. The thiazolidine ring also showed no significant engagement. In contrast, the transfer efficiency of the proton at the stereogenic centre of the open β-lactam ring (NCH) and the benzene sidechain CH2 indicated proximity to the binding side. These data reveal substantial uniformity in the binding poses adopted by the MIL series of antibodies, with binding dominated by the benzene ring. We selected MIL-3 for further biophysical and structural characterisation. We performed surface plasmon resonance (SPR) with a PenG-coated chip, which revealed a modest KD value (5.28 µM; 90% CI: 3.996– 7.37) (**Fig 5d,e**).

To generate a structural understanding of the clonotypic response to penicilloyls, we determined the crystal structure of the PenG-Lys MIL-3 complex at 2.2 Å (**Table S1**). Three Fab molecules were present in the asymmetric unit (H (heavy)/L (light), A/B and C/D). In both H/L and A/B molecules, the phenylacetamide and the thiazolidine moieties are well-defined in the electron density whilst the Cβ, Cδ and Cγ portion of the lysine moiety has weak density, indicating it is less well ordered. The electron density is considerably weaker in A/B than in H/L most likely due to crystal contacts present in H/L (**Fig S7a**). Apart from this region, the interactions between the ligand and the protein are conserved in both FAbs. In the third Fab molecule (C/D), CDRH loops are highly disordered, and no ligand was fitted.

Our analysis focuses on the H/L molecules (**Fig 5f**). The PenG benzene group is deeply buried in a narrow hydrophobic pocket sandwiched on one face of the benzene ring by CDR L loops (Tyr68 L, Tyr110L) and the β-turn formed by CRDH3, mainly the side chain of Ile120H with some involvement of the main chain of Phe124H, on the other face (**Fig 5f ii**., **FigS7b**). The interaction with Tyr68L has strong ρε-stacking character whilst the planes of the rings benzene and Tyr110L are offset 70° and thus hydrophobic in character. The methyl group of Cψ2 Ile120H sits centred above plane of the ring.

The structure does not disclose a simple explanation for the observation from NMR for the interactions of the *p*-hydrogen. The nitrogen of the amide forms a hydrogen bond with the carbonyl of Thr121H whilst the carbonyl oxygen bridges via a water molecule (W1) to the side chain hydroxyl of Tyr110L. The β-turn conformation of CDRH3 is stabilised by hydrogen bonds between residues Thr122H and Arg123H and Tyr 51H and Asp75L.The carboxylate of the thiazolidine moiety makes a bidentate salt bridge to the guanidine group of Arg69H and potentially a salt bridge with His54H (**Fig 5f iii**.); this grouping is likely to powerfully contribute to binding enthalpy. The dimethyl group and thiazolidine ring make van der Waal contacts with Tyr115L and Ile120H while the nitrogen of the ring interacts with a highly coordinated water molecule (W3) bridging CDRs H1, H3 and the lysine linker. The main-chain mimic of the Lys makes five hydrogen bonds with the 310 helix of CDRH2 in H/L (**Fig 5f iv**.) but only one in A/B (**Fig S7c**), consistent with differences in ordering. The main chain atoms of a lysine adduct on a protein would be unable to make these interactions, but torsion angle modification of the lysine side chain would permit the main chain portion (that would be part of the modified protein) to reach outside the binding pocket without any perturbation of the thiazolidine and benzene interactions.

### Evaluation of antibody clones on antibiotic sequestration

The biophysical properties of the antibodies, the structural biology and the physiological concentrations of antibody *in vivo* suggest that the anti-PenG antibodies were unlikely to have a potential drug-sequestering effect. Nevertheless, we evaluated this by developing a model system to test whether PenG-specific antibodies could reduce the bacterial growth-inhibiting function of PenG. Whole antiserum was added to a culture of attenuated, unencapsulated *Streptococcus pneumoniae* (*38*), chosen because of its high sensitivity to PenG (minimum inhibitory concentration (MIC) = 0.01 µg/mL). We conducted two assays: first, we evaluated the effects of anti-HSA- PenG antiserum on PenG sequestration in culture and, second, contrasted PenG kill zones when drug was pre-incubated with high quantities of purified recombinant control or anti-PenG MIL antibodies (**Fig S9**). Neither antiserum nor recombinant MIL antibody significantly inhibited antibiotic action. Collectively, these data are not indicative of antibodies with detectable drug-sequestering or other inhibitory activity, coordinate with the structural and biophysical characterisation.

## Discussion

The elicitation of anti-drug antibodies such as those against PenG may have broad pharmacokinetic and pharmacodynamic implications, including the mediation of drug hypersensitivity (*7*, *14*, *39*, *40*). The complex relationship between chemical instability, pharmacokinetics, immunological factors and possible downstream functional effects of the resulting humoral responses are poorly understood. To address this, we have systematically evaluated the way in which the protein conjugation reactivity of the common β-lactam antibiotic, PenG, drives formation of antigenic complexes *in vitro*, and established that diverse protein carriers are sufficient to propagate a drug-specific IgG response. Using a murine model, we characterised both the clonal B cell and antibody responses revealing clonal restriction and conservation of antibody binding features. Collectively, our data provide a model for how adduct formation and immune engagement triggers the humoral response, phenomena that may inform both the study of allergy and potentially provide a rational approach to predict and even mitigate anti-penicillin antibody responses. Our data demonstrate that the production of PenG- specific antibodies is regulated at two distinct levels: 1) the covalent formation of protein adducts via lysine-amide formation is influenced by reaction medium and time, and 2) immune engagement, including innate recruitment—such as via adjuvant which *in vivo* would be mimicked by bacterially-elicited inflammation—and T cell help. These factors ultimately determine the overall probability and magnitude of a downstream B cell response against PenG.

In a human context, most patients undergoing standard courses of PenG to treat bacterial infection exhibit anti-penicilloyl IgG antibodies thereafter (*6*, *41*, *42*). These data appear initially incongruent with our murine data since animals given formulations of free penicillin either intravenously or in drinking water failed to exhibit a specific response. However, it is noteworthy that these selective effects may be attributed to dosing (since humans are given as much as 50 mg/kg every 4–6 h of PenG via an intravenous line) (*43*, *44*) and/or a higher murine cardiac output and drug clearance rate that will result in shorter drug half-life (*30*, *31*). This is supported by the serological data gathered from mice given both static pre-formed serum-PenG immunisations and intramuscular slow-release PenG-Ben.

It should be noted that these experiments were not designed to explicitly evaluate allergic outcomes in mice; for example, isotype-switching to IgE and the downstream mast cell-mediated and other modes of allergic reactivity, is determined by both genetic factors in the form of atopic predisposition to IgE production, and environmental factors (*45*, *46*). However, the functional effect of the antibody response may not only be a feature of the Fc effector function or isotype—for example, whether type I versus type II hypersensitivity is imparted—but also the inherent binding mode and biophysical properties of the antibody. Our data show that the murine B cell repertoire responds to the drug adduct with a dominant clonotypic family that primarily engages the side-chain constituent, benzylacetamide, and the carboxylate group of the thiazolidine, as evidenced by our complementary uSTA (*37*) and x-ray structural characterisation. This is consistent with previous mapping studies of murine (*47*, *48*), rabbit (*23*) and human (*49*) antibody responses in which side-chain reactivity appears prominent. Moreover, the biophysical features of these antibodies, with monovalent

Fab KD in the low–mid µM range and relatively fast koff, coupled with lower concentrations *in vivo*, appear together to discount even partial drug inhibitory effects.

Our approach, applied here to an archetypal inhibitor with a covalent bond-forming mode of action, creates a potential roadmap for how to instruct the chemical, pharmacological and immunological factors governing whether a B cell response can be mounted against β-lactam and other potentially protein-reactive therapeutics such as proton pump inhibitors (*5*). We describe here in a mouse that PenG adducts are recognised by a unique B cell clonotype, with notably narrow VH gene utilisation and the identification of key motifs that interact with the antibiotic sidechain motif. Having thus identified a dominant clonotype against the PenG adduct, this excitingly suggests a workflow that could be used as a model to reverse engineer a PenG analogue that fails to engage the MIL clonotype, potentially reflecting a low/no allergenic alternative against a murine germline and that now instructs a proof-of-principle for germline-informed drug design in humans.

## Methods

### Ex vivo modification of protein with β-lactam antibiotics

Carrier proteins (HEL, OVA, HSA, BSA and MSA) were purchased commercially (Merck) and dissolved in 0.1 M NaCO3 (pH = 8, unless otherwise indicated) and concentration was adjusted to 1 mg/mL. β-lactam was added to a molar ratio of 1:100 per lysine residue of carrier protein. The mixture was rotated end-to-end at 25°C overnight and dialysed into PBS.

### Denaturing MS

Reversed-phase chromatography was performed in-line prior to mass spectrometry using an Agilent 1100 HPLC system (Agilent Technologies inc. – Palo Alto, CA, USA). Concentrated protein samples were diluted to 0.02 mg/ml in 0.1% formic acid and 50 µl was injected on to a 2.1 mm x 12.5 mm Zorbax 5um 300SB-C3 guard column housed in a column oven set at 40°C. The solvent system used consisted of 0.1% formic acid in ultra-high purity water (Millipore) (solvent A) and 0.1 % formic acid in methanol (LC-MS grade, Chromasolve) (solvent B). Chromatography was performed as follows: Initial conditions were 90% A and 10% B and a flow rate of 1.0 mL/min. After 15 s at 10% B, a two-stage linear gradient from 10% B to 80% B was applied, over 45 s and then from 80% B to 95% B over 3 s. Elution then proceeded isocratically at 95% B for 1 mins 12 s followed by equilibration at initial conditions for a further 45 s. Protein intact mass was determined using a 1969 MSD-ToF electrospray ionisation orthogonal time-of-flight mass spectrometer (Agilent Technologies Inc. – Palo Alto, CA, USA). The instrument was configured with the standard ESI source and operated in positive ion mode. The ion source was operated with the capillary voltage at 4000 V, nebulizer pressure at 60 psig, drying gas at 350°C and drying gas flow rate at 12 L/min. The instrument ion optic voltages were as follows: fragmentor 250 V, skimmer 60 V and octopole RF 250 V. Obtained MS spectra were processed and deconvoluted using the Agilent MassHunter Qualitative Analysis (B.07.00) software.

#### LC-MS/MS

Approximately 5 µg protein was reduced, loaded and run on an SDS-PAGE. Gel bands were excised and washed sequentially with HPLC grade water followed by 1:1 (v/v) MeCN/H2O. Gel bands were dried (via vacuum centrifuge), treated with 10mM dithiothreitol (DTT) in 100mM NH4HCO3 and incubated for 45 minutes at 56°C with shaking. DTT was removed and 55mM iodoacetamide (in 100mM NH4HCO3) was added and incubated for 30 minutes in the dark. All liquid was removed and gels were washed with 100mM NH4HCO3/MeCN as above. Gels were dried and 12.5 ng/µL trypsin was added separately and incubated overnight at 37°C. Samples were then washed and peptides were extracted and pooled with sequential washes with 5% (v/v) formic acid (FA) in H2O and MeCN. Dried samples were reconstituted in 2% MeCN 0.05% trifluoroacetic acid and run by LC-MS.

Samples were analysed using an Ultimate 3000 UHPLC coupled to an Orbitrap Q Exactive mass spectrometer (Thermo Fisher Scientific). Peptides were loaded onto a 75 µm × 2 cm pre-column and separated on a 75 µm × 15 cm Pepmap C18 analytical column (Thermo Fisher Scientific). Buffer A was 0.1% FA in H2O and buffer B was 0.1% FA in 80% MeCN with 20% H2O. A 40-minute linear gradient (0% to 40% buffer B) was used. A universal HCD identification method was used. Data was collected in data-dependent acquisition mode with a mass range 375 to 1500 m/z and at a resolution of 70,000. For MS/MS scans, stepped HCD normalized energy was set to 27, 30 and 33% with orbitrap detection at a resolution of 35,000.

Raw data was first searched using the FragPipe (v19.1) (*50*) Open Search pipeline to determine the modified mass shift caused by PenG conjugation. Approximately 2.3% of peptide spectral matches (PSM) had an unannotated mass shift of 334.099 Da. Next, to determine site specificity and occupancy, raw data was searched using Proteome Discoverer (3.0.0.757). In-house curated FASTA databases were used. Digestion enzyme was set to trypsin with a maximum of 2 miss cleavages. A 10 ppm precursor mass tolerance and 0.6 Da fragment mass tolerance were allowed. Oxidation (+15.995 Da) of methionine and PenG conjugation (+334.099 Da) of lysines and protein N-termini were set to dynamic modifications. Carbamidomethylation (+57.021 Da) of cysteines was set as a static modification. Target false discovery rate (FDR) for peptide spectrum matches, peptide and protein identification was set to 1%. To approximate site specific occupancy of PenG conjugation the total number of PSMs of peptides containing a specific lysine site in a PenG modified state were expressed as a percentage compared to the total number of PSMs containing the given lysine in both the modified and unmodified state.

### Mice and immunisation formulations

Wild-type, specific pathogen-free, sex-matched, 6–8-week-old C57BL/6 mice were purchased from Charles River. Animals were monitored daily and were provided standard chow and water *ad libitum*. Immunisation formulations and schedules are outlined in the results. Mice were bled periodically from the tail vein. Animals were sacrificed via a rising CO2 gradient and subsequent cervical dislocation schedule 1 procedure. All experiments were conducted under the approved licenses, consistent with national and University guidelines.

#### ELISA

Serum samples were serially diluted and transferred onto an antigen-coated and blocked SpectraPlate-96 (PerkinElmer) plate. Binding was detected with an anti-mouse IgG-HRP (STAR120P, Bio-Rad). ELISAs were developed using 1-Step-Ultra TMB ELISA substrate (Life Technologies), terminating the reaction with 0.5 M H2SO4. For competition ELISAs, serial dilution of soluble ligands as preincubated with antiserum at the pre-determined EC50 concentration for 1 h. The antisera and ligand mixtures were subsequently transferred onto the antigen-coated and blocked plates and ELISA conducted, as previously outlined. Cytokine ELISAs were performed using commercially available kits (Life Technologies), screening supernatant from antigen-restimulated splenocytes.

Optical densities were measured at 450 and 570 nm on a Spectramax M5 plate reader (Molecular Devices). After background subtraction, logistic dose-response curves were fitted in GraphPad Prism. Endpoint titres were determined as the point at which the best-fit curve reached an OD450-570 value of 0.01, a value which was always > 2 standard deviations above background.

### B cell sorting

Penicilloyl-specific B cells were isolated using antigen probes. Gp120-PenG was modified with an NHS-esterified AF647 dye, as per the manufacturer’s instructions (Life Technologies). To improve true antigen-specific cell sorting efficiency, a negative backbone-specific probe was synthesised, wherein biotinylated gp120 was tetramerised with PE-conjugated strepdavidin (Biolegend).

Single cell suspensions were stained with LIVE/DEAD Fixable Blue and Fc receptors blocked. Surface staining was performed using anti-mouse F4/80-PE (1:200, BM8, Biolegend), anti-mouse Gr-1 (1:200, RB6-8C5, Biolegend), anti-mouse CD3-PE (1:200, 17A2, Biolegend), anti-mouse CD4-PE (1:200, RM4-5, Biolegend), anti-mouse CD8-PE (1:200, RPA-T8, Biolegend), anti-mouse B220-eFluor450 (1:100, RA3-6B2, BD Biosciences), anti-mouse IgD-AF700 (1:200, 11-26c.2a, Biolegend), anti-mouse IgM-PE/Cy7 (1:200, R6-60.2, BD Biosciences), anti-mouse IgG1-FITC (1:200, A85-1, BD Biosciences), anti-mouse IgG2a/2b-FITC (1:200, R2-40, BD Bioscience) and antigen probes (10 μg/mL). Cells were stained on ice for 1 h, washed and stored on a BD FACSAriaFusion (BD Biosciences). Single cells were sorted into MicroAmp Optical 96-well PCR plates (Life Technologies), isolating LIVE/DEAD^-^DUMP^-^B220^+^IgD^-^gp120^-^ gp120-PenG^+^ events. Cells were sorted directly into 5 µL if 1X TCK buffer supplemented with 1% 2-ME and stored at -80°C until use.

### Variable region cloning and antibody expression

B cell receptor variable regions were recovered, as previously described (*25*). Briefly, RNA was captured on RNAClean XP beads (Beckman Coulter) and washed with 70% ethanol. RNA was eluted and cDNA was synthesised using SuperScript III (Life Technologies) with random primers (Life Technologies). VH and VK regions were recovered (*35*) and Q5 polymerase (New England Bioscience), sequencing the amplicons via Sanger. VH amplicon sequences were used to determine B cell clonality.

To validate that the sequences were specific to the penicilloyl adducts, antibodies were recombinantly expressed. The VH/VK amplicons were incorporated into expression vectors: vector-overlapping adapters were incorporated via PCR (*34*), and the V regions were inserted into pre-cut recombinant Fab expression (*36*) vector via Gibson reaction (New England Bioscience). Vector products were transiently transfected into HEK 293Freestyle cells and Fab was purified from supernatant using Ni-NTA resin.

### Immunogenetic analyses

Analyses were performed using VH regions. Sequences were aligned to the murine reference genome using the Immunogenetics Information System (IMGT; https://www.imgt.org/IMGT_vquest/input), as described previously (*25*). Sequence outputs of poor quality or those unproductive were excluded from our analyses. Alignments of CDRs were visualised using WebLogo (*51*). Clonal lineages were evaluated using GCTree (*52*).

#### SPR

SPR was performed using a Biacore T200 instrument. Details of chip design, synthesis and testing, refer to **Document S1**. Fab binding was evaluated by sequentially injecting serial dilutions at a flow rate of 10 µL/minute.

### uSTA

Samples for uSTA were prepared by buffer-exchange of purified fab fragments with D2O PBS using Amicon 30K MWCO. All NMR experiments were conducted on Bruker Avance Neo 600 MHz spectrometer at 25°C equipped with QCIF cryoprobe and a SampleJet, running TopSpin 4.2.0. The uSTA experiments were recorded using stddiffesgp.2 pulse sequence as a pseudo 3D. The saturation times were as follows: (0.1, 0.3, 0.5, 0.7, 0.9, 1.1, 1.3, 1.5, 1.7, 1.9, 2, 2.5, 3, 3.5, 4 and 5 s). On-resonance spectra were recorded with Gaussian excitation pulse at 9 ppm at off-resonance recorded with Gaussian excitation pulse at 37 ppm. Other frequencies were also tested, however they showed direct excitation of the ligand (**Figure S1**). Experiments were recorded with 32768 number of points and sweep width of 16.01 ppm. Data were processed using nmrPipe workflow, integrated into the uSTA data collection. Transfer efficiencies were calculated by subtracting the on-resonance spectra from the off-resonance spectra and normalising this to the 1D off-resonance spectra according to the equation: Transfer Efficiency=STD/1D, where STD = off-resonance - on-resonance. The highest intensity peak (H-20, NCHCOOH, thiazolidine ring) was used as a reference peak. The heatmaps were generating by encoding the percentage transfer efficiency onto the chemical structure of PenG. Interpolation of transfer efficiencies in the heatmaps on carbon, sulphur, oxygen, and nitrogen was performed following the known 1/d6 dependency of dipolar interactions, as described previously (*37*).

*X- ray crystallography*

MIL-3 Fab was loaded onto a gel filtration Superdex 200 column 10/30 (GE Healthcare) in 10 mM Tris-HCl, pH 7.5, 150 mM NaCl. Co-crystals appeared at 20 C after a week from a hanging drop of 0.1 μL of protein solution (15 mg/mL with 2.5 mM PenG-Lys with 0.1 μL of reservoir solution containing 20% (w/v) PEG 6000, 0.1 M MES pH 6, 0.2 M ammonium chloride in vapor diffusion with reservoir. Crystals were frozen with the same solution containing 16% glycerol. Data were collected at the Diamond light source oxfordshire (beamlines I04). Data were processed with XIA2 (*53*–*56*). Structure has been solved by molecular replacement using PHASER and pdb file 7bh8 for VL, CH and CL domains and 7n09 for VH domain. The structure was builded with Autobuild program, refined with REFINE of PHENIX with NCS restraints (*57*) and adjusted with COOT (*58*). Coordinates and topologies of ligands were generated by AceDRG (*59*).

### Microbiological assays

The attenuated, unencapsulated lab strain *Streptococcus pneumoniae* D39 (Δ*cps2A’*- Δ*cps2H’*) (*38*) was routinely grown in tryptic soy broth (TSB) (BD Biosciences) at 37°C (standing incubation) in a 5% CO2 atmosphere.

Microbroth dilutions of S*. pneumoniae* D39 revealed a PenG MIC of 0.01 μg/mL (that is, the lowest antibiotic concentration that prevented bacterial growth) (data not shown). For the liquid antibiotic rescue assay, 2 ng of PenG in 10 μL PBS were pre-incubated with 10 μL of antisera for 2 h in a flat-bottom 96-well plate. 180 μL of exponentially growing *S. pneumoniae* cells (OD600 = 0.2) were added and incubated overnight (such that the final PenG concentration was 0.01 μg/mL). The following day, bacterial growth was measured using a Spectramax M5 plate reader (Molecular Devices), evaluating the OD600 nm as a proxy for bacterial density.

For disk diffusion assays, 0.1 μg of PenG and mAb (1:50 molar ratio) were spotted onto paper disks. The disk was placed onto a blood agar plate (Merck), carrying a 5 ml nutrient soft agar overlay with 200 μL exponentially growing *S. pneumoniae* cells (OD600 = 0.2). Plates were incubated overnight. The following day, the kill zone diameter was manually measured. PenG-only, PenG-raised antibody clones were tested, as well as an irrelevant HIV-1-specific antibodies were tested.

## Data and statistics

Flow cytometry data was evaluated using FlowJo V.10.8.2 for Mac. Statistical analyses were conducted in either GraphPad Prism V.10.0.1 or in RStudio V.4.1. Statistical test details are provided in the results, figures and associated figure legends.

## Acknowledgments

We thank The Sir William Dunn School of Pathology flow cytometry facility, SPR facility and animal house staff. We extend our gratitude to the Rosetrees Trust, who have supported this work through the Interdisciplinary Award (ID2020/100023). We are also grateful for the funding provided by the Wellcome Trust (224212/Z/21/Z). Additionally, we thank the Wellcome Trust (grant ref: 095872/Z/10/Z) and the Engineering and Physical Sciences Research Council (grant ref: EP/R029849/1) for the instrumental upgrades of the 600-mHz and 950-MHz NMR spectrometers, as well as support from the University of Oxford Institutional Strategic Support Fund, the John Fell Fund, and the Edward Penley Abraham Cephalosporin Fund. AJB is supported by ERC grant (101002859). For the purpose of Open Access, the author has applied a CC BY public copyright license to any Author Accepted Manuscript version arising from this submission. The Chemistry theme at the Rosalind Franklin Institute is sustained by the EPSRC (V011359/1 (P)). We would like to thank the Membrane Protein Laboratory at Diamond Light Source (funded by Wellcome Trust grant 223727/Z/21/Z) for help and support. CMK is supported by an EPA Cephalosporin Junior Research Fellowship from Linacre College Oxford. LPD is supported by the Clarendon Fund, and QJS is a Jenner Vaccine Institute Investigator and James Martin School Senior Fellow.

## Author contributions

Conceptualization of project: LPD, BGD, QJS

Methodology: LPD, LM, GS, VL, AT, CJB, SAB, CMK, MS, WBS, AJB, JN, BGD, QJS

Investigation: LPD, LM, GS, VL, AT, CJB, SAB, CMK, WBS

Funding acquisition: MS, WBS, AJB, JN, BGD, QJS

Project administration: MS, WBS, AJB, JN, BGD, QJS

Supervision: MS, WBS, AJB, JN, BGD, QJS

Writing – original draft: LPD

Writing – review & editing: LPD, BGD, QJS

## Supplementary Document 1

### 1. Synthesis of penicillin-lysine ligand 1

#### General method

All reagents were purchased from commercial sources and were used without further purification unless noted. Dry solvents for reactions were purchased from Fisher Scientific, where the solvent product series of ‘[Solvent name], 99.9 % Extra Dry, AcroSeal^TM^’ were applied for all reactions. Following abbreviations are used: PE = petroleum ether (b.p. 40 – 60 °C), EtOAc = ethyl acetate, THF = tetrahydrofuran. Thin Layer Chromatography (TLC) was carried out using Merck aluminium-backed sheets coated with Kieselgel 60-F254 silica gel. Visualization of the reaction components was achieved using UV fluorescence (254 nm) and/or by charring with an acidified p-anisaldehyde solution in ethanol or an acidified cerium ammonium molybdate (CAM) solution in water. Organic solvents were evaporated under reduced pressure. Lysine-penG product was purified by a Teledyne Combiflash Nextgen 300 system with a 50 g HP C18 Gold cartridge. The gradient for Combiflash purification is: 5% B at 0 CV, 5% B at 1 CV, 50% B at 11 CV, 100% B at 14 CV, 100% B at 16 CV (Solvent A = Milli Q water, solvent B = acetonitrile). Proton Nuclear Magnetic Resonance (^1^H NMR) spectra were recorded on a Bruker AVB400 (400 MHz), or AV700 (700 MHz) spectrometers, and the chemical shifts are referenced to residual CHCl3 (7.26 ppm, CDCl3), CHD2OD (3.30 ppm, CD3OD), HDO (4.79 ppm, D2O). Carbon Nuclear Magnetic Resonance (13C NMR) spectra were recorded on a Bruker AVB400 (100 MHz), or AV700 (175 MHz) spectrometers and are proton decoupled, and the chemical shifts are referenced to CDCl3 (77.16 ppm) or CD3OD (49.0 ppm). Assignments of NMR spectra were based on two-dimensional experiments (^1^H-^1^H COSY, DEPT-135, HSQC, and HMBC) if required. Reported splitting patterns are abbreviated as s = singlet, d = doublet, t = triplet, q = quartet, p = pentet, hept = heptet, m = multiplet, br = broad, app = apparent. Low Resolution Mass Spectra (LRMS) were recorded on a Waters ACQUITY QDA mass spectrometer using electrospray ionization (ESI). High Resolution Mass Spectra (HRMS) were recorded on a Waters Xevo-G2 QTof spectrometer using electrospray ionization (ESI), m/z values are reported in Daltons.

### Synthesis of (2R,4S)-2-((R)-2-(((S)-5-acetamido-5-carboxypentyl)amino)-2-oxo-1-(2-phenylacetamido)ethyl)-5,5-dimethylthiazolidine-4-carboxylic acid (1)

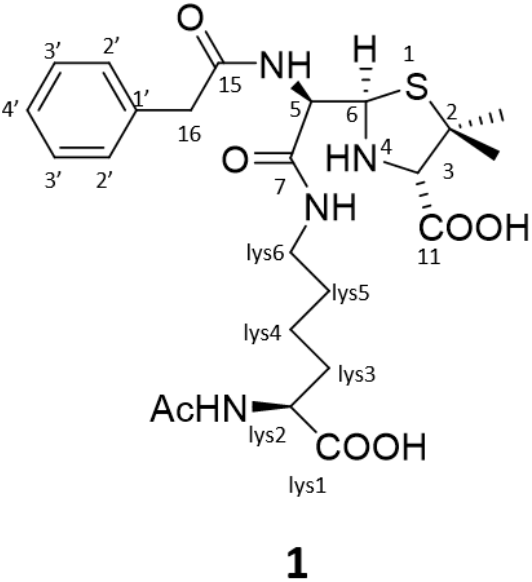

Penicillin G sodium salt (20 mg, 0.056 mM) and Nα-*Acetyl*-L-*lysine (30 mg, 0.159 mM)* was dissolved in 0.5 mL sodium carbonate buffer (0.1 M, pH = 11). The solution was stirred at room temperature overnight. When TLC showed the completion of reaction, the entire reaction mixture solution was injected into a C18 combiflash reverse phase purification system (see general method for more details, it is important not to use acidified mobile phase). Fractions containing the product was combined and passed through a small column of Amberlite IR-120 resin (Na^+^ form). The solvent was removed under vacuum to afford the product as a white solid (22 mg, 75%, the actual reaction yield is almost 100%, we discarded those mixed fractions for purity which sacrificed the final recovery yield).

Characterisation: TLC: Rf = 0.5 (2:2:1 Ethylacetate: Isopropyl alcohol: water). 1H NMR (400 MHz, D2O) δ 7.32 (dd, *J* = 8.0, 6.4 Hz, 2H, H3’), 7.31 – 7.20 (m, 3H, H2’ & H4’), 4.82 (d, *J* = 9.3 Hz, 1H, H6), 4.20 (d, *J* = 9.3 Hz, 1H, H5), 3.99 (dd, *J* = 9.2, 4.5 Hz, 1H, Lys-H2), 3.64 – 3.51 (m, 2H, H16), 3.40 (s, 1H, H3), 3.15 – 2.99 (m, 2H, Lys-H6), 1.94 (s, 3H, Lys-AcNH), 1.68 (dtd, *J* = 13.2, 7.7, 4.6 Hz, 1H, Lys-H3), 1.51 (s, 3H Me2 + Lys-H5a), 1.37 (dtt, *J* = 62.6, 13.4, 6.6 Hz, 1H, Lys-H5), 1.23 (q, *J* = 7.9 Hz, 2H, Lys-H4), 1.17 (s, 3H, Me1).13C NMR (101 MHz, D2O) δ 179.51, 174.97, 174.44, 173.54, 171.10, 160.89, 134.89, 129.13, 128.91, 127.31, 74.37, 64.62, 59.35, 59.25, 55.27, 42.02, 39.04, 31.16, 27.90, 27.15, 26.73, 22.71, 22.00. HRMS: Calculated [M] = 522.2148 Da, found [M + H+] = 523.4766 Da.

#### Spectrums for compound 1

Low resolution MS (left) and example TLC for the reaction (right, 2 hours’ time point)

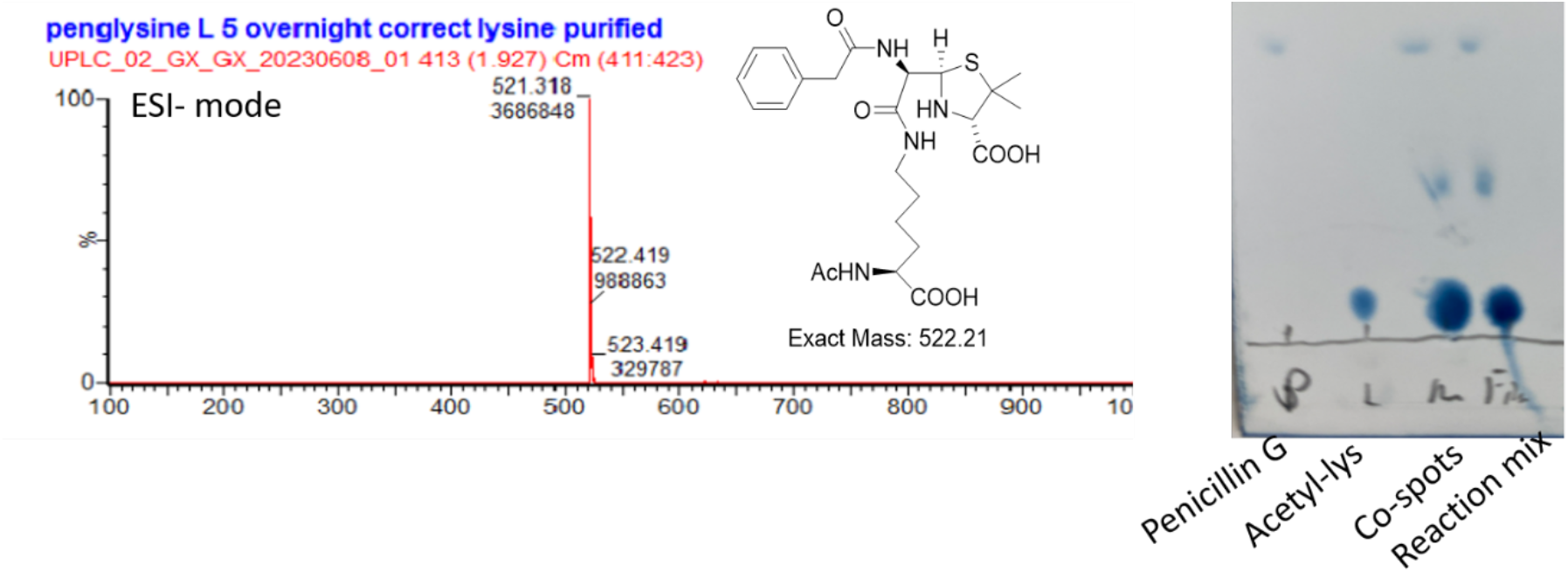

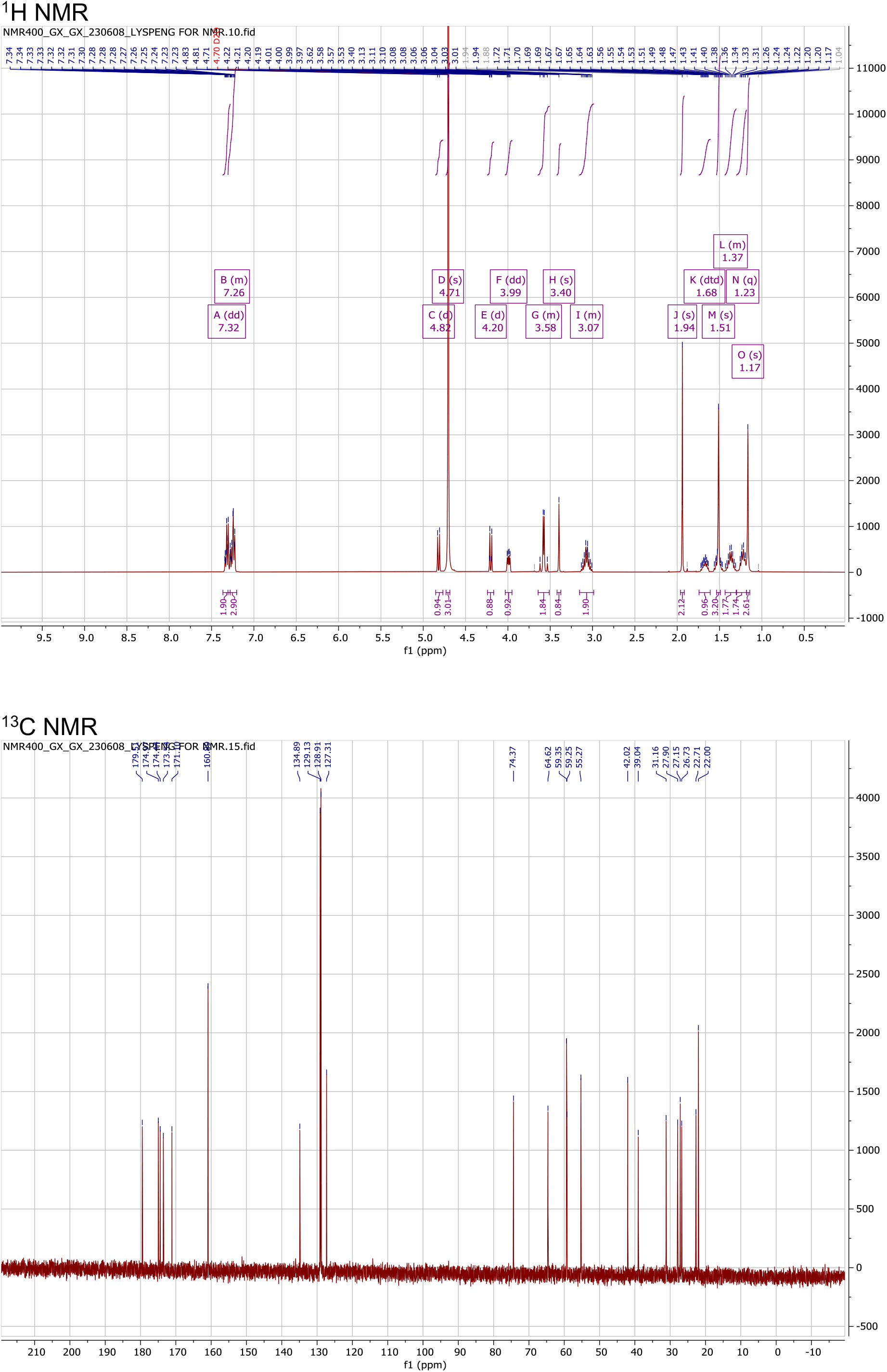

HMBC confirmed the ring opening-linkage between lysine and penicillin G, due to HMBC spectra showed LysH6 coupled with C7 of penicillin G.

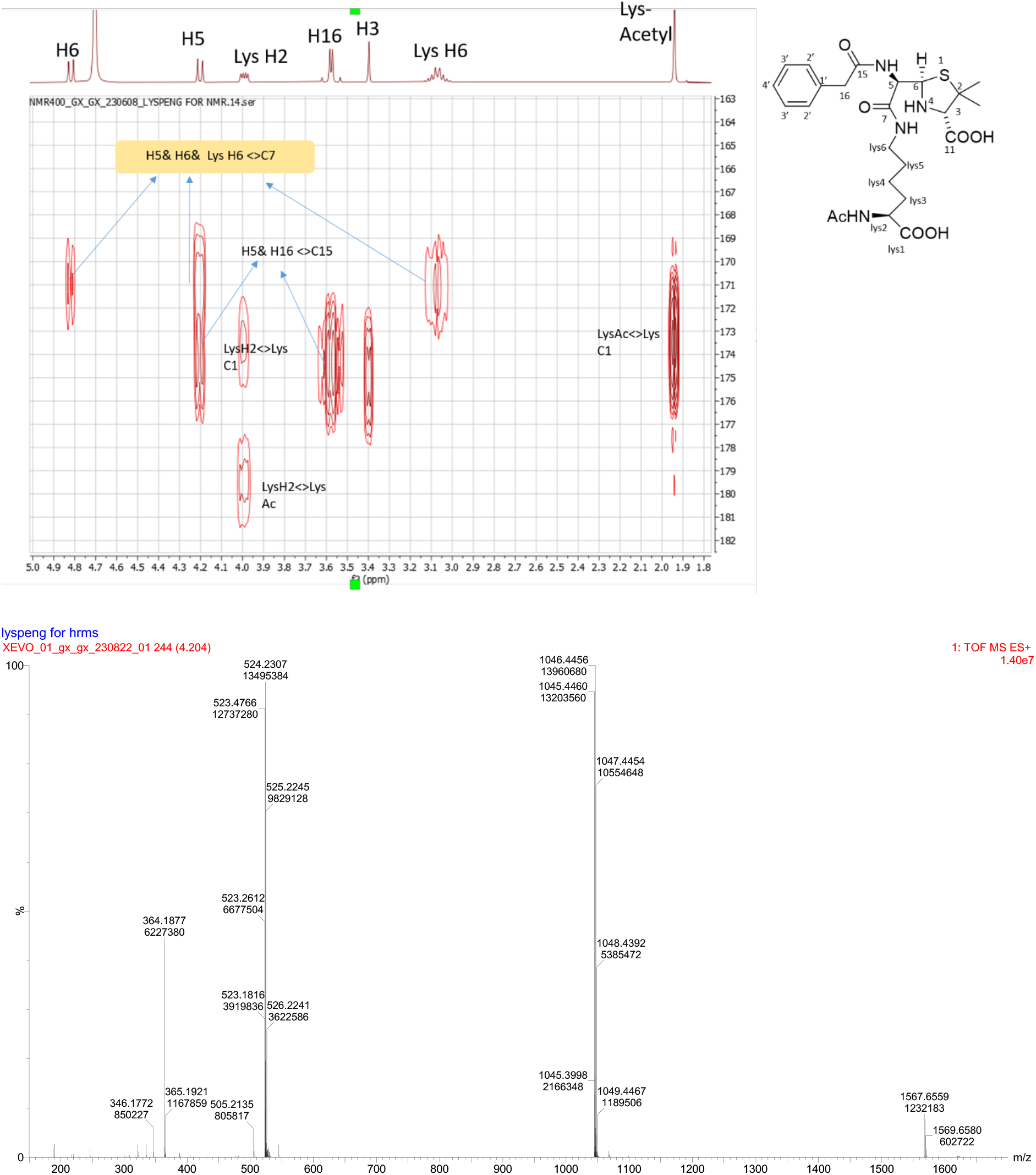

HRMS for compound 1: Calculated [M] = 522.2148 Da, found [M+H^+^] = 523.4766.

### 2. Preparation of penG-lysine SPR chip

#### PenG-lysine stability test

Ligand stability is critical to the success of ligand modification on SPR chips. Due to there is no literature which have reported the stability of compound 1, we carried out a small stability test to find the best storage condition for our penG-lysine SPR chip. Experimental design: 3 mg of compound 1 was dissolved in 700 uL D2O PBS and the pH was adjusted to 2, 8 and 11. Such 3 samples was loaded to 400 mHz NMR machine and NMR spectra by different time point was recorded.

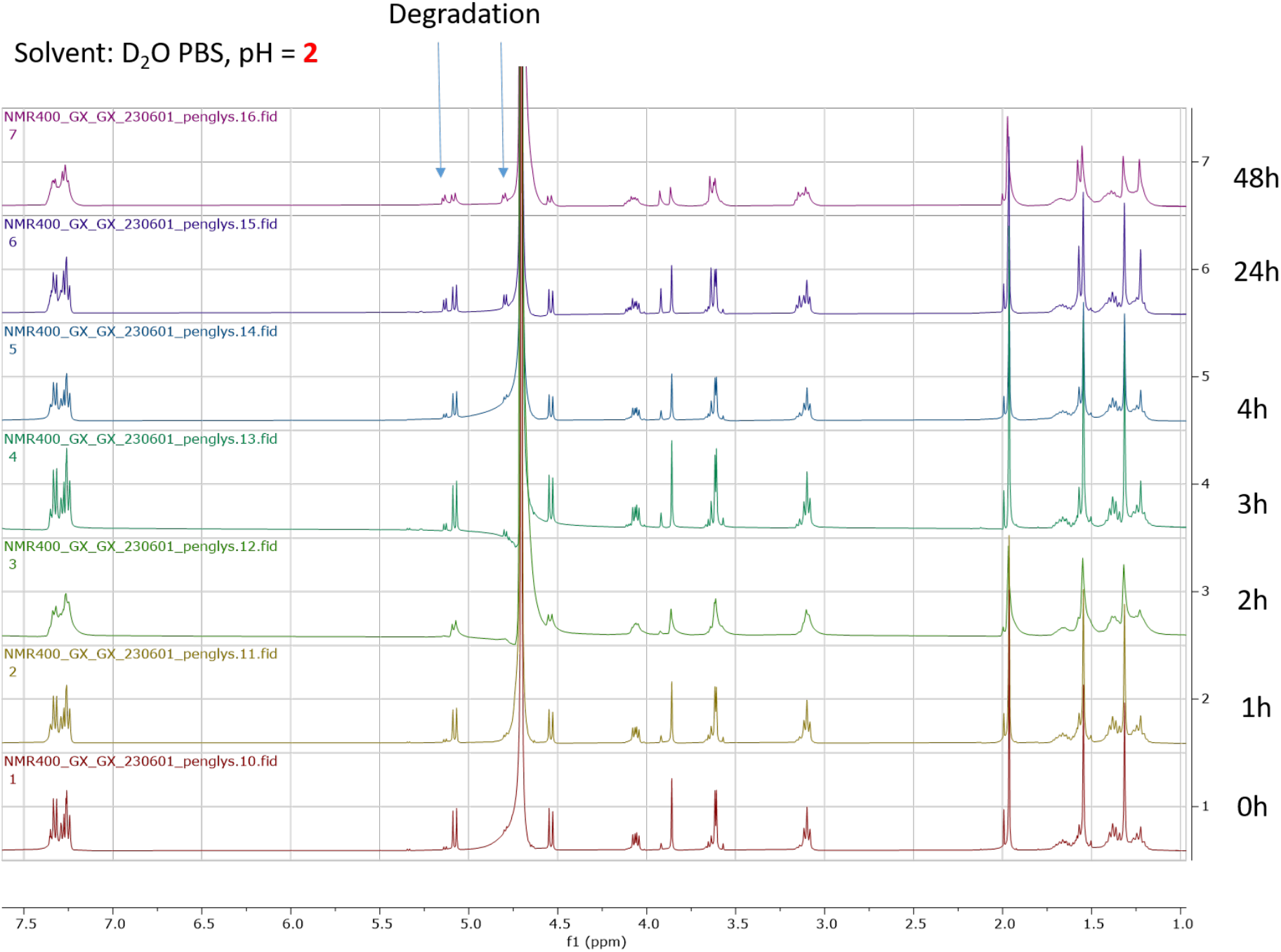

**Figure 1:**
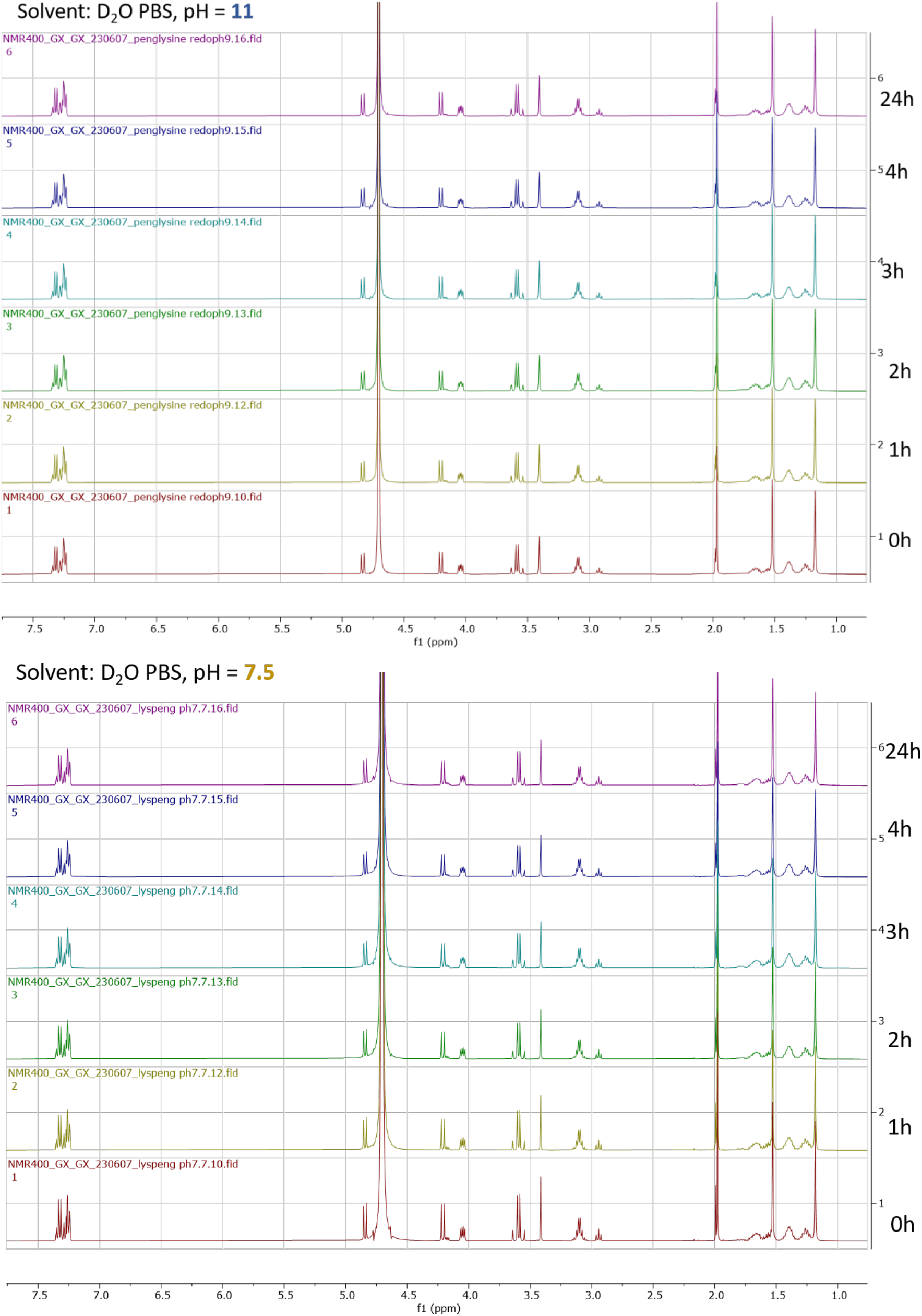
Stability test of penG-lysine (1). Samples were prepared in D2O PBS and pH was adjusted to relevant value showed in the figure. We only observed degradation when pH = 2. Compound **1** in pH = 11 & 7.5 D2O PBS is stable. Therefore, we decided to use PBS for the storage of penicillin G SPR chip.

#### Preparation of penicillin G SPR chip

**Scheme 1:**
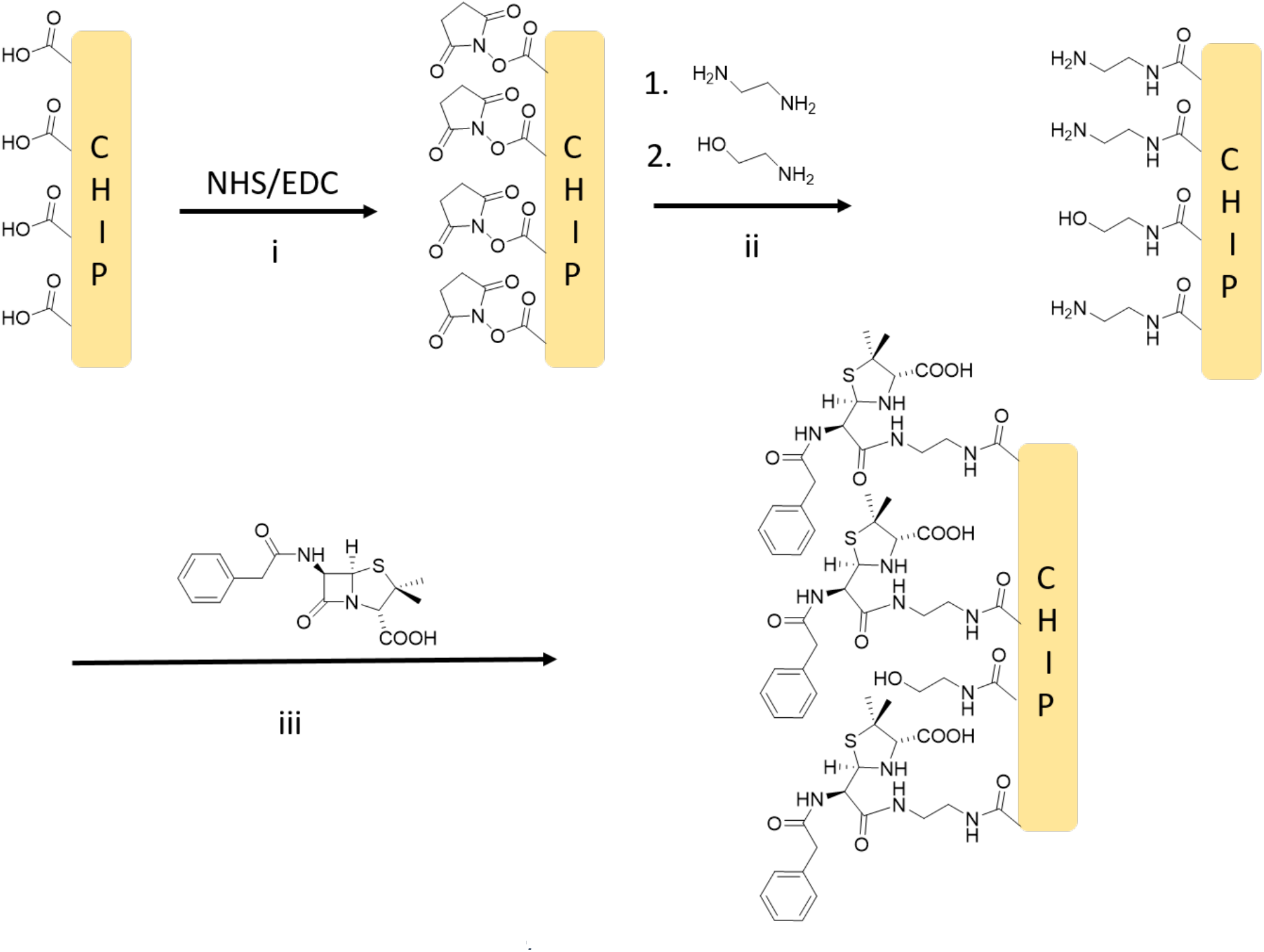
Preparation of penicillin G SPR chip.

Immobilisation of penicillin-G onto carboxymethylated SPR sensor chip. A carboxymethylated CM5 sensor chip was activated with N-hydroxysuuccinimide (NHS) and N-ethyl-N′-(3-dimethylaminopropyl)carbodiimide hydrochloride (EDC•HCl) and subsequently coupled to ethylenediamine followed by blocking of unreacted esters with ethanolamine. Penicillin G was dissolved in sodium carbonate buffer (pH = 11) and was then injected into the flow cell, where it reacts with the surface bound free amines to mimic the conformation of penicillin G-lysine.

Specifically, in step i, a CM5 chip activated with NHS (50 mM) and EDC (200 mM) 10 min at 10 μL/min. In step ii, ethylenediamine (1 M) was injected to functionalise chip with free amine groups for 7 min at 10 μL/min. Then ethanolamine (1 M) was injected over 10 min to block any unreacted NHS activated esters at 10 μL/min. In step iii, penicillin G sodium (112 mM) was injected over 150 min at 5 μL/mL followed by washing with PBS buffer (wash steps with buffer between each step and during sensor equilibration). The prepared chip was stored in PBS (pH = 8) to prevent from dry out and degradation.

### 3. Preparation of protein-penG antigen

The beta-lactam antibiotic’s ring-opening by nucleophilic attack from protein lysine residue has been well studied in many literatures, therefore, we decided to start with a literature method [1] where such reaction was done in 0.1 M Na2CO3 pH = 11 buffer. We selected hen egg lysozyme (HEL) as a model protein for this reaction. We also prepared samples with different pH to define the optimum pH.

#### Experimental

0.1 M Na2CO3 bsuffer, range from pH = 11, 10, 9, 8, 7.8, 7.6, 7.4, 7.2, 7.0, 6.8 was prepared. For each pH entries was added: 0.2 mg HEL commercial HEL protein powder and 200 eq. per-lysine (1200 eq. of HEL protein) penicillin G sodium salt. The final concentration of protein is 1 mg/mL. The solution was stirred overnight at room temperature and the number of modification was visualized by protein LC-MS. The molecular weight of unmodified HEL is 14306 Da.

#### Results

##### Discussion and conclusion (Figure 2)

**Figure 2:**
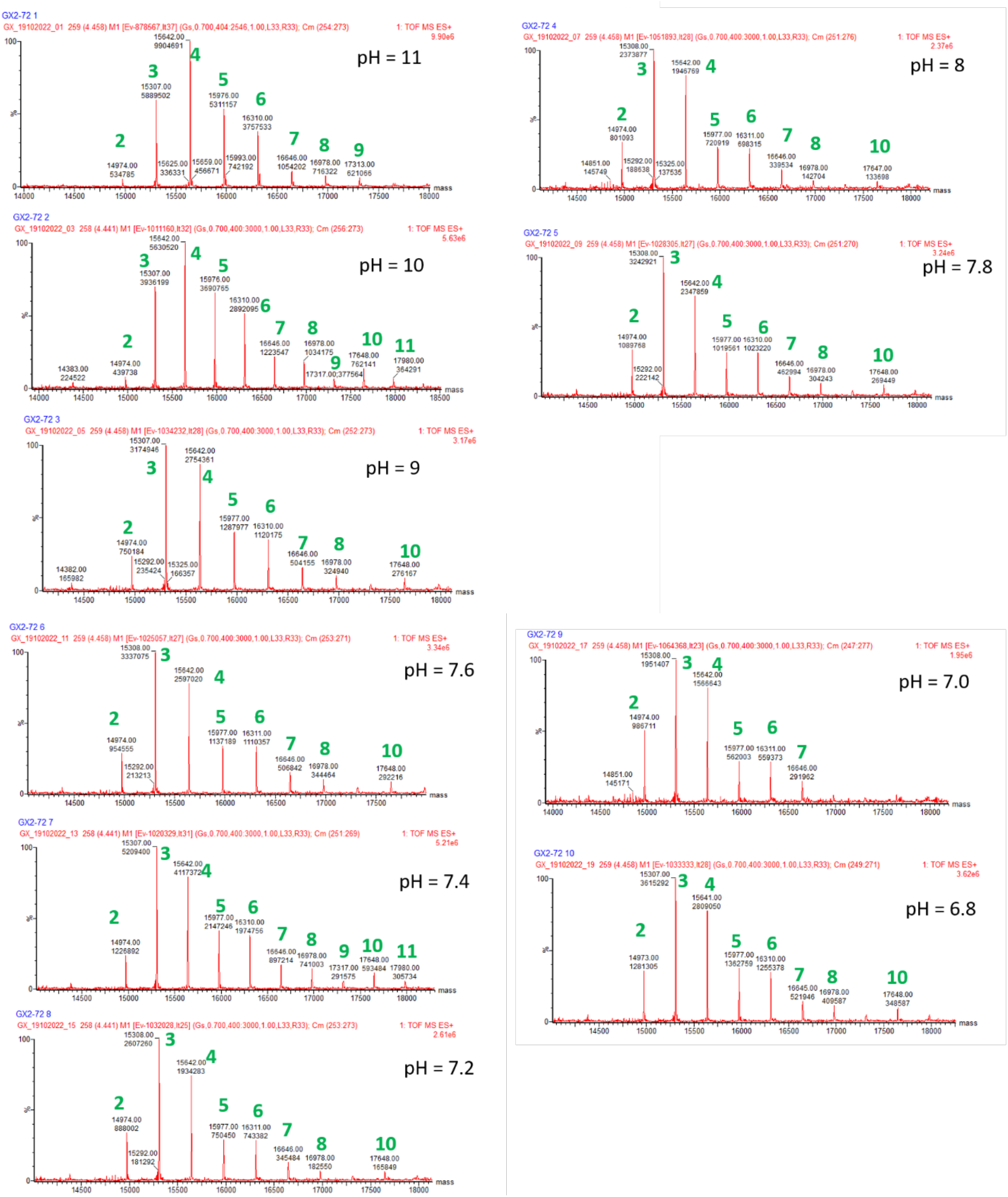
Penicillin G modified HEL at different pH in carbonated buffer. The green number represents the number of penicillin molecule loaded to the HEL protein, e.g. **2** modification means: 14306 Da (Mass of naked HEL protein) + 334 Da (Mass of penicillin G) *2 = 14874 Da.

Sodium carbonate buffer is preferred for penicillin G-protein complex preparation, as it resulted in at least 2 modifications product up to 11 modifications. Changing pH of sodium carbonate buffer does not significantly affect the modification results. Moreover, due to HEL protein only have 6 lysine and 1 N-terminal, the maximum number of modification is expected to be 7. However, we did see modification number > 7, which indicated that another non-lysine type of protein residue was also modified at this specific reaction condition. We finally chose pH = 11 for all the same reaction mention in this paper.

#### Additional test for non-carbonated buffers

We repeated the penicillin-HEL reaction described in previous section but changed the reaction buffer from Na2CO3 to Milli Q water (Figure 3, a) and commercial PBS buffer (pH = 7.2, Figure 3, b) to investigate the effect of carbonate buffer.

**Figure 3:**
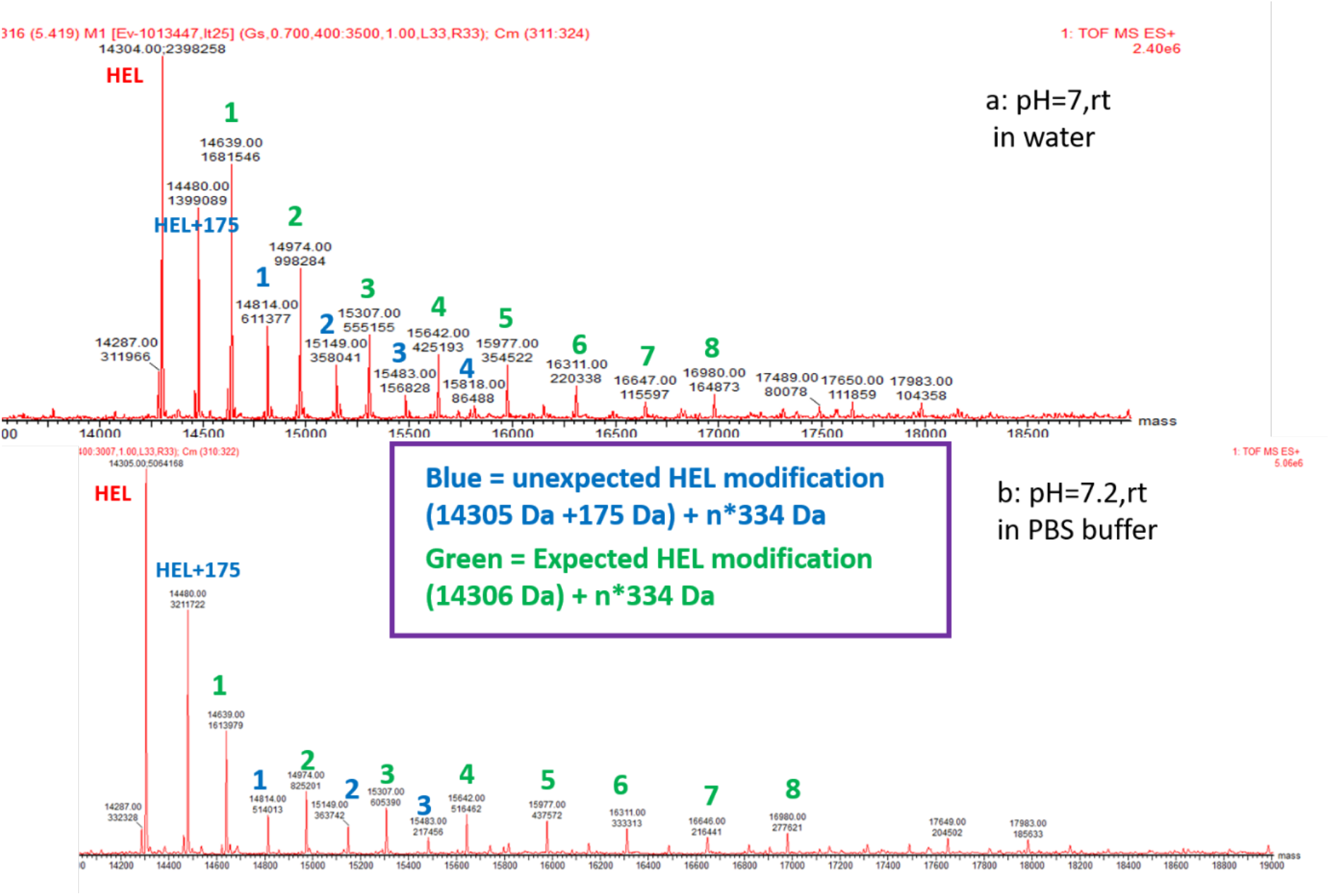
Penicillin G modified HEL in Milli Q water and PBS. The green numbers represent the number of penicillin molecule loaded to the HEL protein, e.g. **2** modification means: 14306 Da (Mass of naked HEL protein) + 334 Da (Mass of penicillin G) \***2** = 14874 Da. Blue numbers represent the unexpected HEL modification from the initial HEL + 175 Da modification. E.g. **2** modification means 14480 Da (14305 Da + 175 Da, naked HEL + unknown modification 175 Da) + 334 Da (Mass of penicillin G* **2**).

The reaction was processed much slower in water and PBS due to a significant amount of unreacted HEL protein was remained after an overnight period of incubation (Figure 3, peak 14305 Da). Surprisingly, we observed a group of patterned and unexpected peaks, where the naked HEL protein was firstly modified by an unknown adduct mass of 175 Da, forming a new peak of 14480 Da (= naked HEL 14035 Da + 175 Da). Then, the new-formed 14480 Da protein was added the whole number times of mass of penicillin G, giving a unique set of peaks added to the spectrum. As a result, the synthesis of penicillin-G antigen should be restricted in carbonated buffer to ensure the product specificity.

---

**Figure S1:**
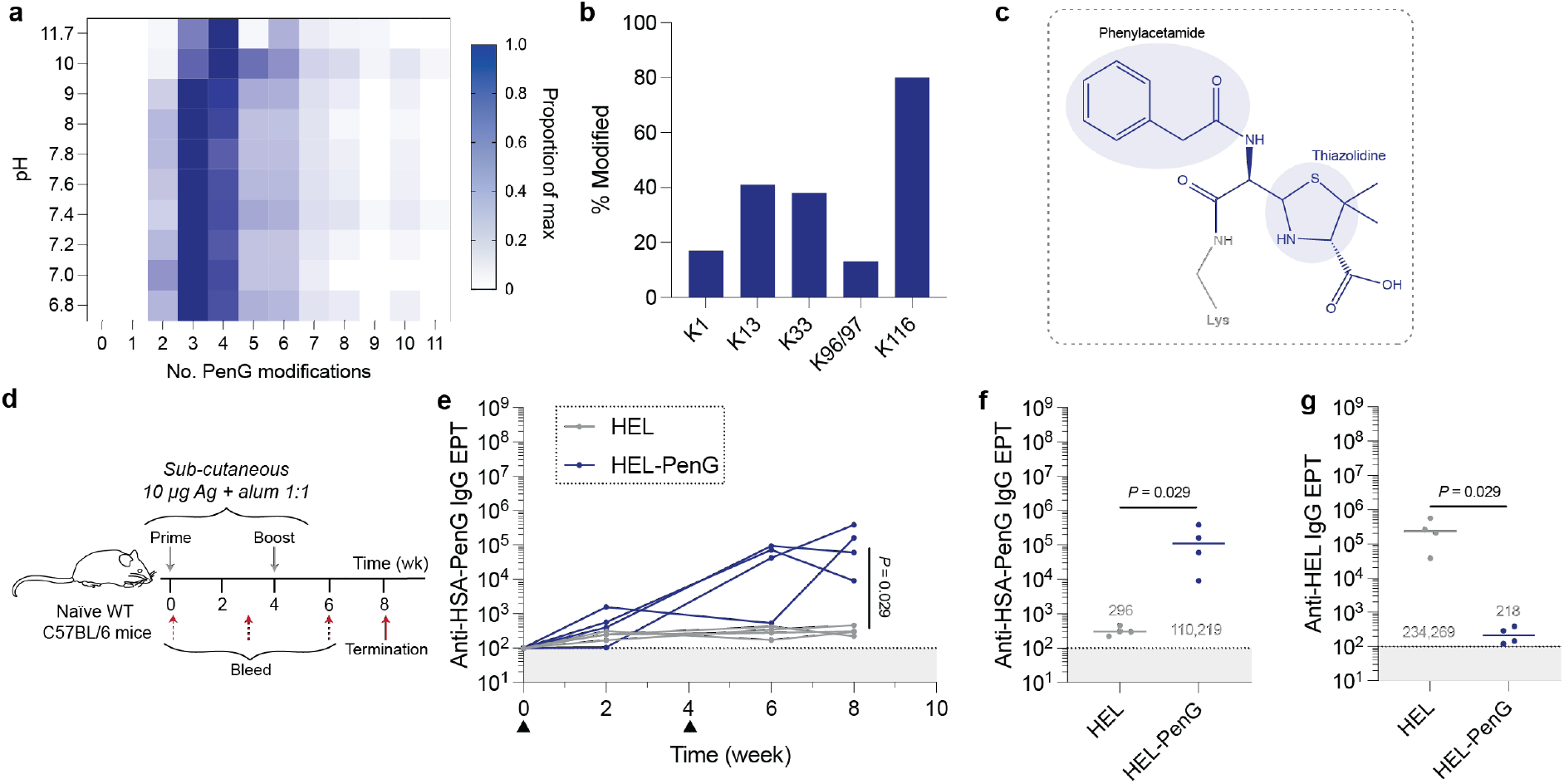
Chemical characterisation of HEL-PenG and its immunogenicity in mice. **(a)** The PenG adduct occupancy was tested with respect to buffer condition, evaluating the relative number of adducts via mass spectrometry. **(b)** Site-specific occupancy of PenG adducts on HEL-PenG (modified at pH = 8.0). **(c)** Inferred PenG-Lys adduct. **(d)** Immunisation schedule. Sex-matched WT 6-week-old C57BL/6 mice were immunised with 10 µg/mL HEL or HEL-PenG in alum. IgG antibody titres against the PenG adduct were evaluated by screening cross-reactivity against HSA-PenG. This was conducted both **(e)** longitudinally and **(f)** at the terminal timepoint. **(g)** The terminal protein backbone-specific response was also determined. Dots represent values from a single animal. Groups were compared via Mann-Whitney tests.

**Figure S2:**
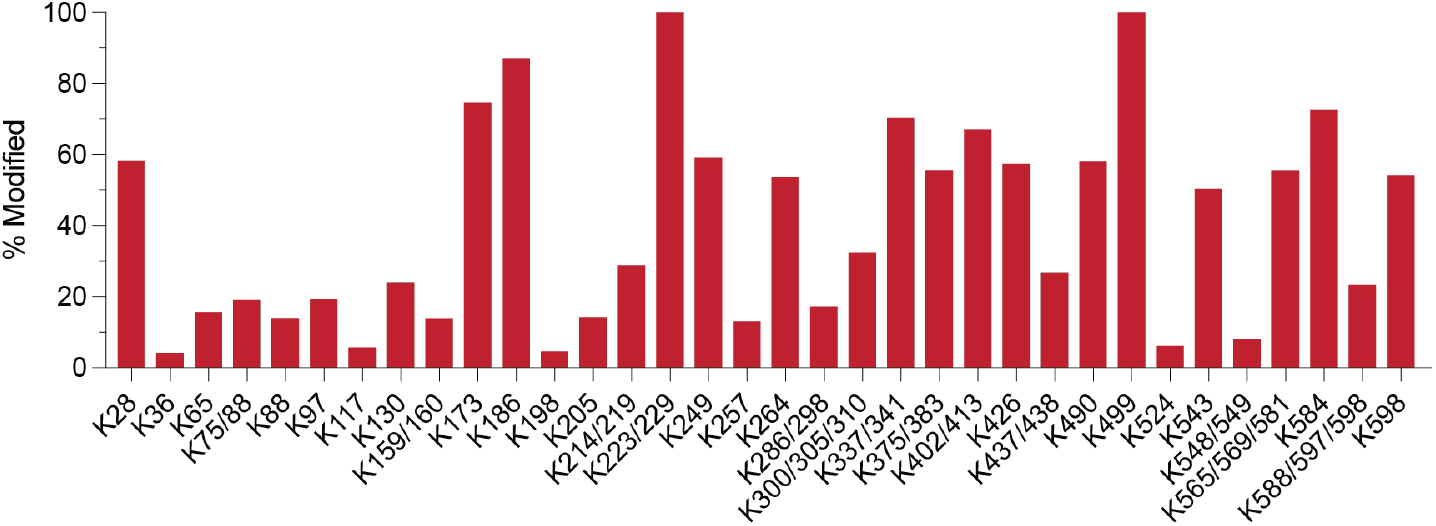
PenG occupancy on HSA. Site-specific occupancy of PenG adducts on MSA-PenG (modified in 0.1 M Na2CO3, pH = 8.0). Determined by tryptic digestion and subsequent evaluation via LC-MS/MS.

**Figure S3:**
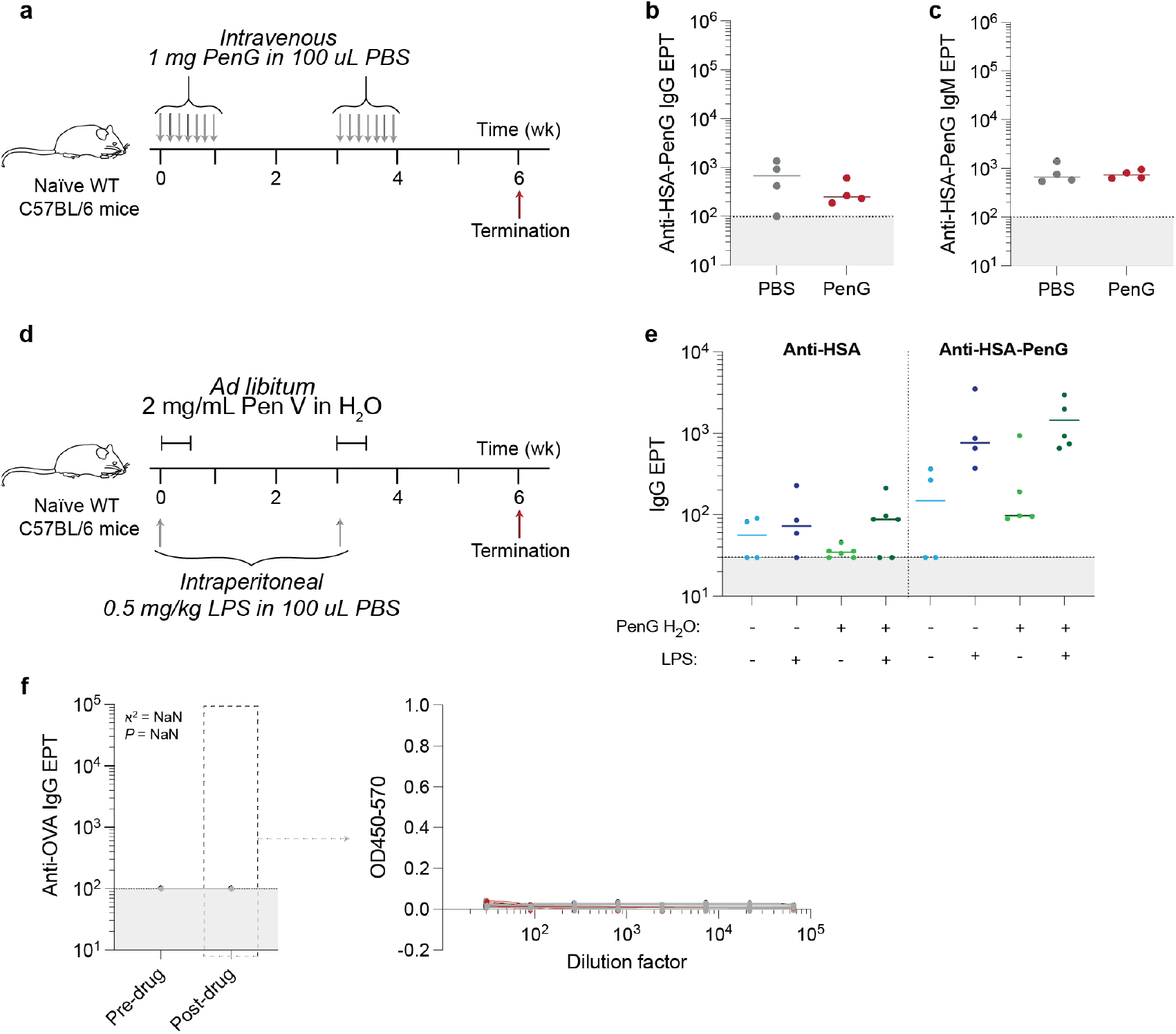
Serological responses following the administration of free PenG/PenV. **(a)** Mice received two week-long courses of PenG intravenously. **(b)** Terminal IgG and **(c)** IgM EPTs were determined against HSA-PenG. **(d)** PenV in water at 2 mg/mL was given to animals *ad libidum*. Some mice revied intraperitoneal immunisations of LPS. **(e)** IgG EPTs against HSA and HSA- PenG were evaluated. **(f)** Anti-OVA EPTs of mice given PenG-Ben intramuscularly.

**Figure S4:**
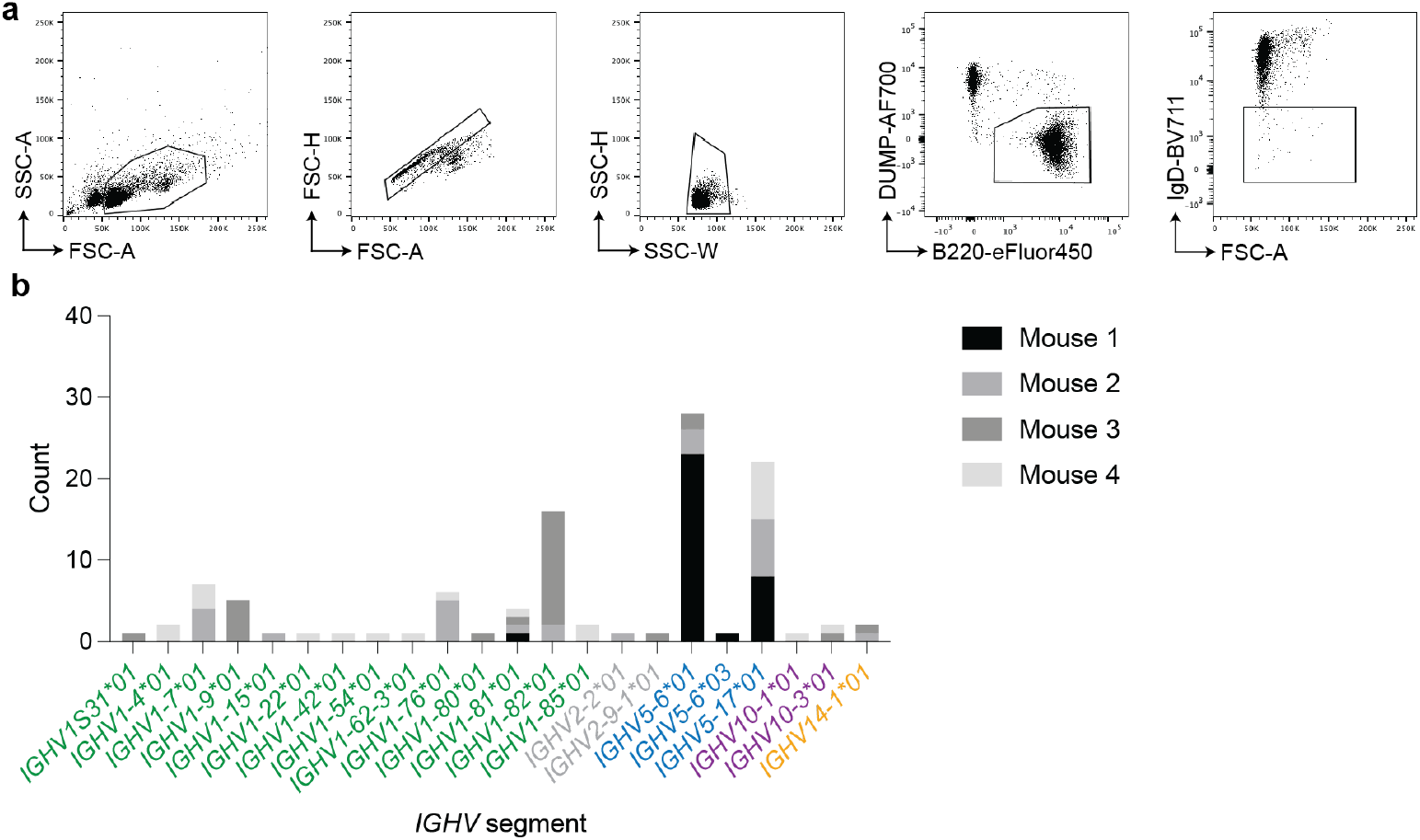
V_H_ gene segments in PenG-specific B cells. **(a)** Flow cytometry pre-gating strategy for mouse IgD^-^ B cells. **(b)** VH gene segments utilized among B cells sorted as PenG-specific. Data pooled from the four mice sequenced.

**Figure S5:**
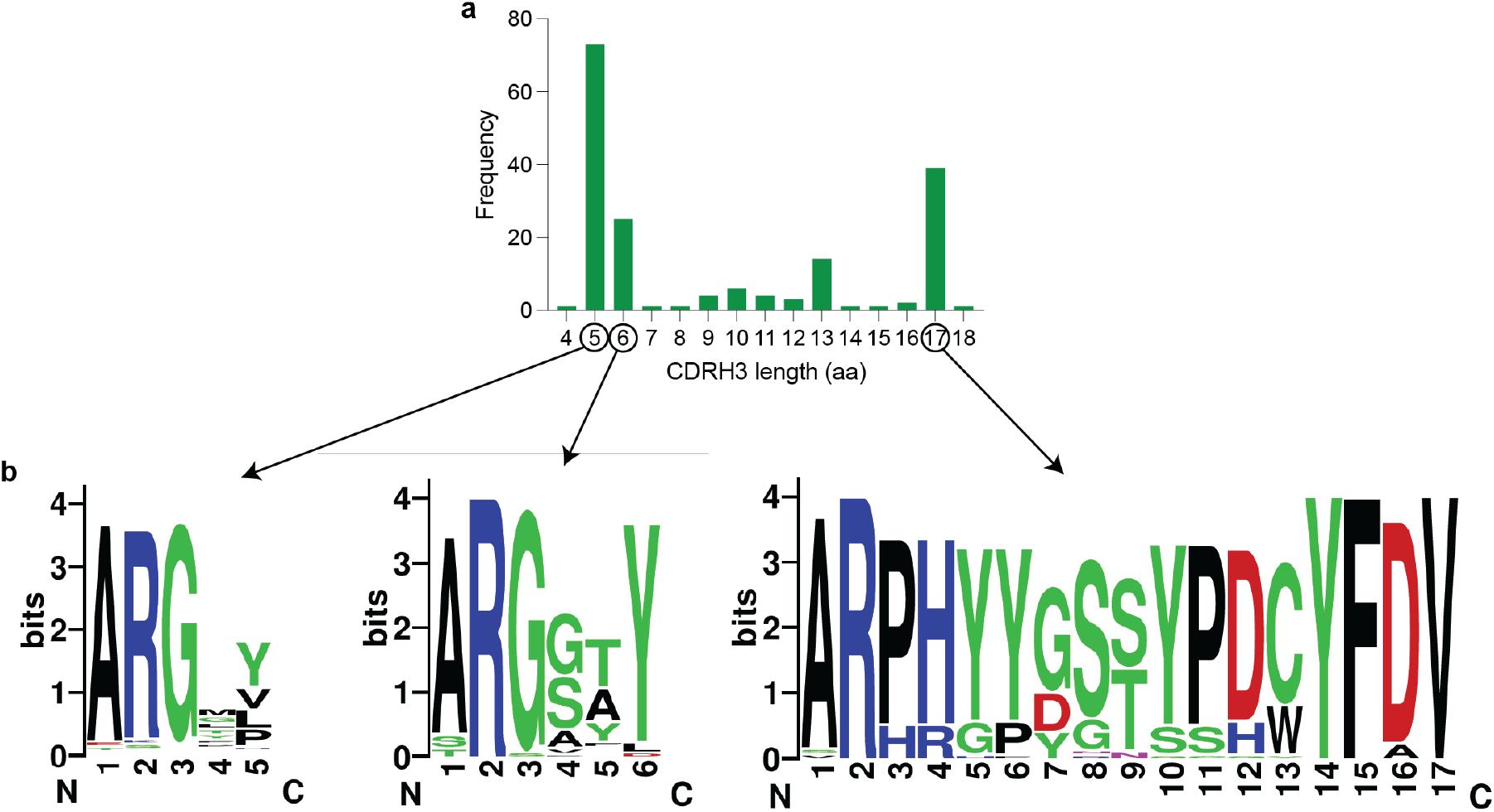
Analysis of CDRH3 in PenG-specific B cell clones. **(a)** Length of CDRH3s of PenG-specific B cells. **(b)** Logo plots of the CDRH3 amino acid sequence from the three most common lengths.

**Figure S6:**
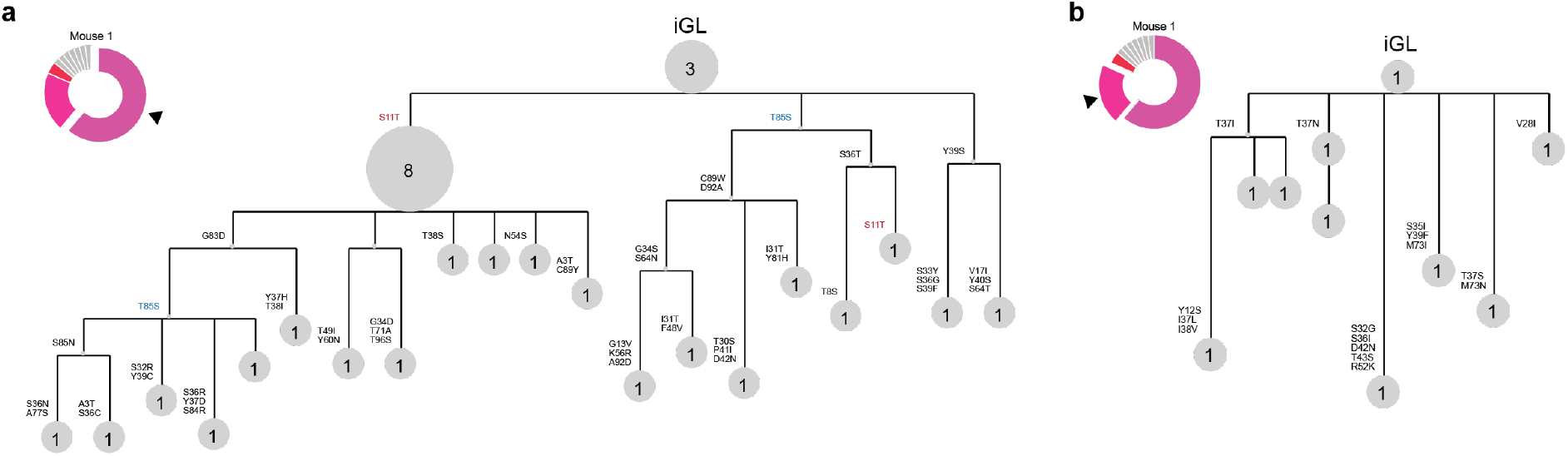
Germinal center trees of large families isolated from Mouse 1. **(a,b)** Germinal center trees of large clonal families isolated from Mouse 1. Number and size of the nodes represents the number of B cells isolated. Amino acid residue mutations are marked.

**Figure S7:**
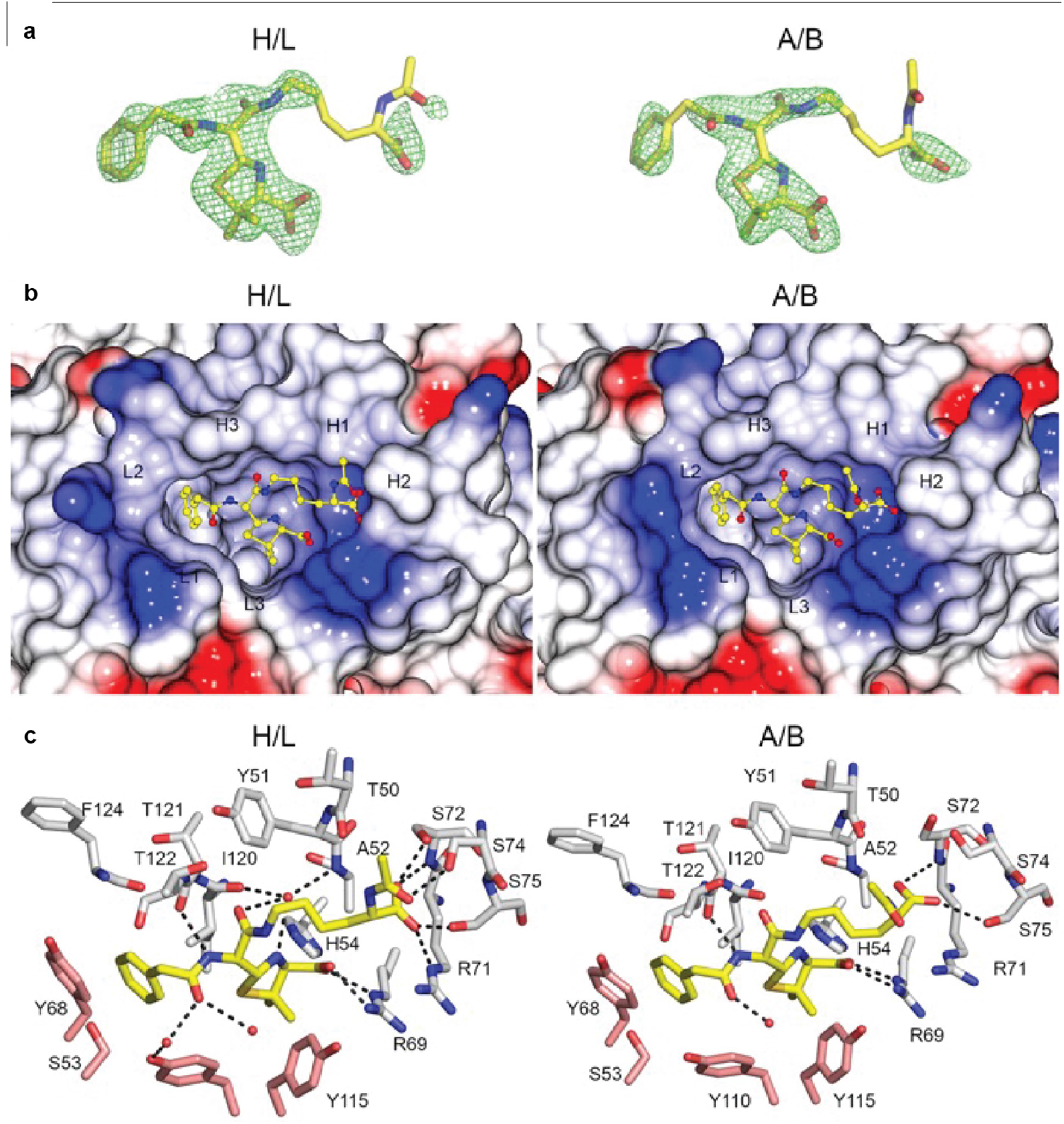
MIL-3 in complex with PenG-Lys x-ray structure. **(a)** FO–FC electron density omit map at 3 σ around PenG-Lys molecule. PenG-Lys is shown as sticks with carbon atoms coloured in yellow, nitrogen in dark blue and oxygen in red. **(b)** Surface of the binding side of MIL-3/ PenG-Lys complex structure. The surface of BMIL-3 is colored by electrostatic charges calculated in CCP4MG (red for negative potential, white for neutral and blue for positive). PenG-Lys is shown as sticks with the carbon in yellow. CDR loops have been labelled**. (c)** Binding site of PenG-Lys. Residues within 4.0 Å of the polysaccharide are displayed and hydrogen bonds are shown as black broken lines. PenG-Lys is shown as sticks with carbon atoms coloured in yellow. Carbon atoms of residues of chain H and L are in white and salmon, respectively.

**Figure S8:**
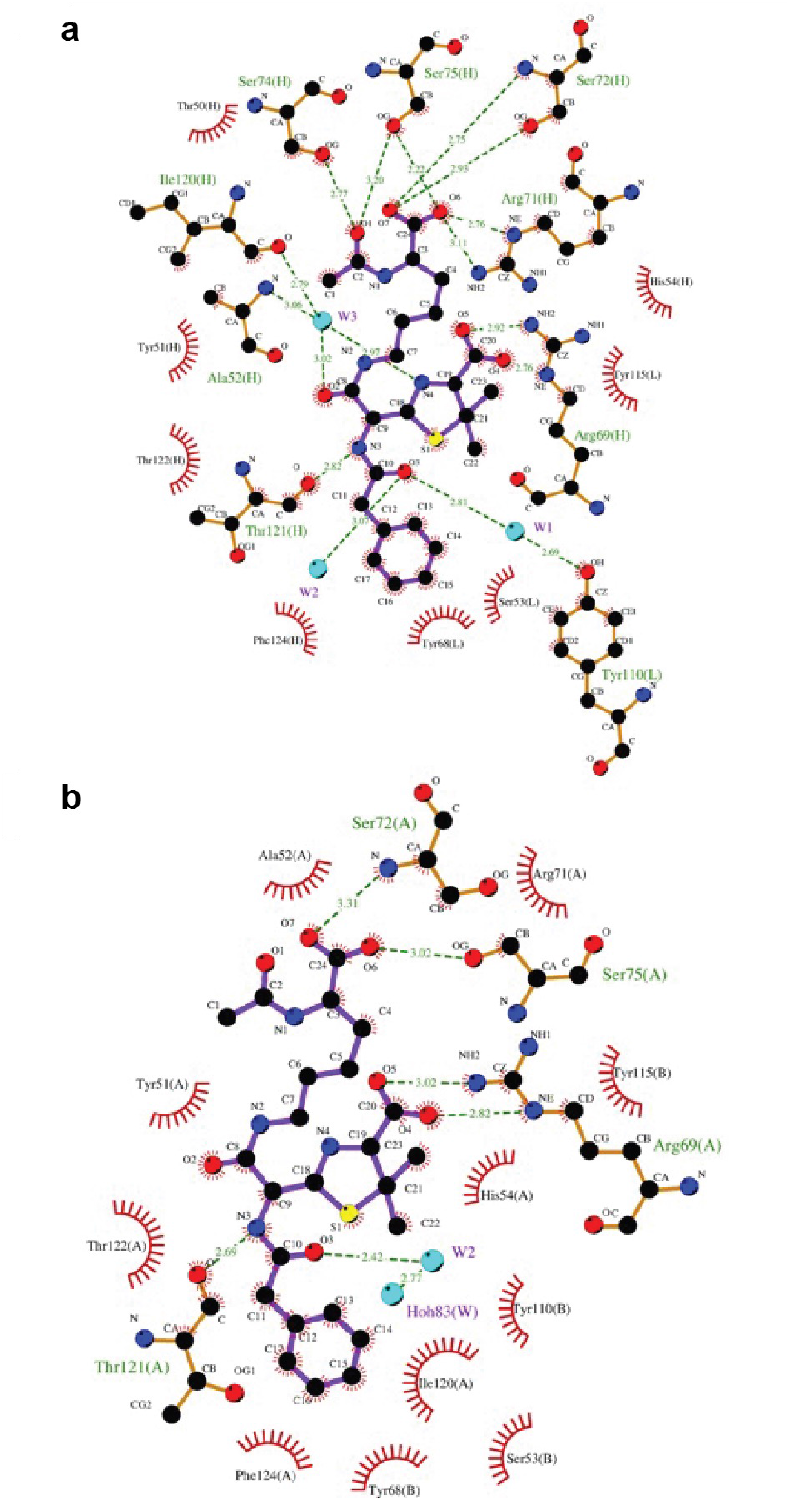
MIL-3 with PenG-Lys ligplot diagrams. Ligplot diagrams illustrating BAR-1/Lys–C(NH)NH-GM3g interactions for chain H/L **(a)** and A/B **(b)**. Covalent bonds of the polysaccharide and the protein residues are in purple and brown sticks, respectively. Hydrogen bonds are represented by green dashed lines and hydrophobic contacts are shown as red semi-circles with radiating spokes.

**Figure S9:**
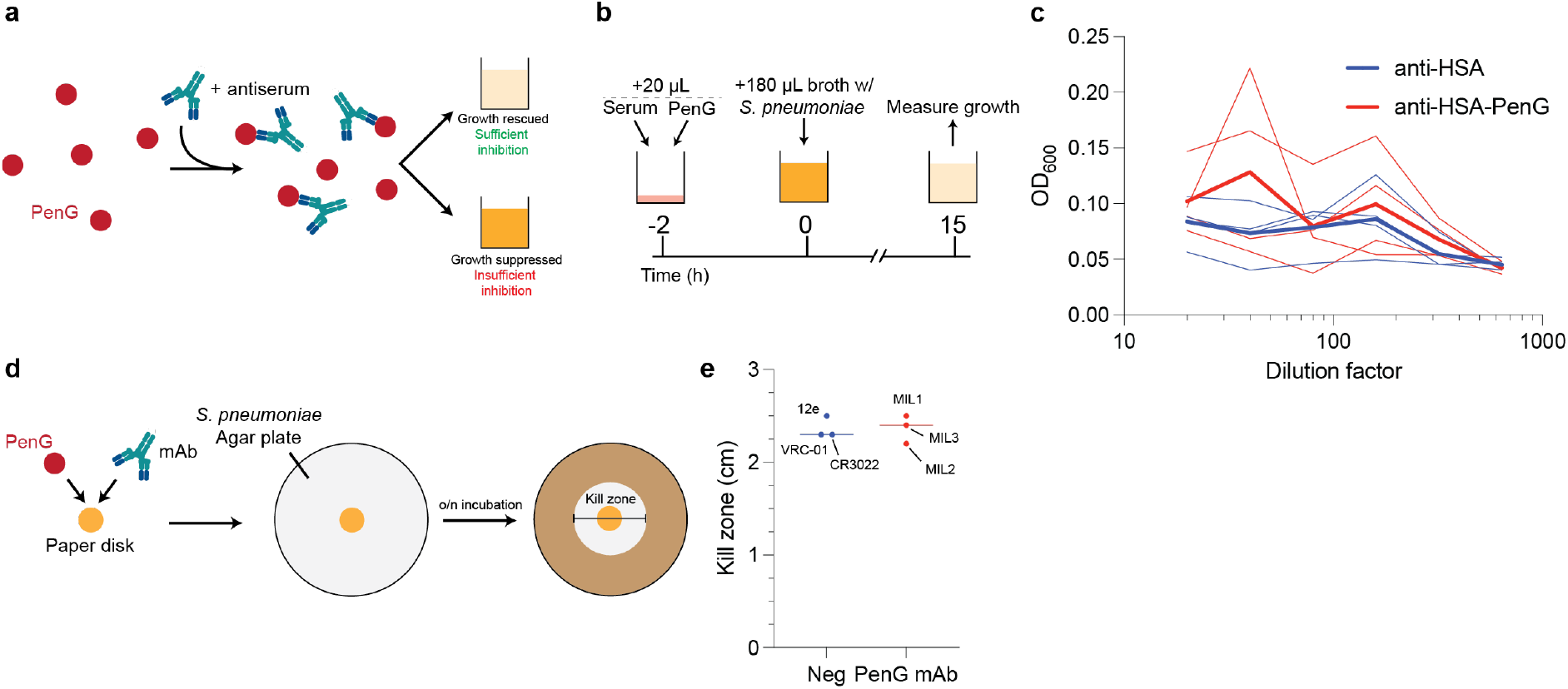
Determining the drug-sequestering effects of the PenG-specific antibody response. **(a)** Methodological concept for liquid antibiotic rescue assay. **(b)** Culture strategy. **(c)** Growth response to pre-incubated antiserum and PenG. **(d,e)** Disk diffusion assay.

**Table S1:** Crystallographic data and refinement statistics.

| 8QXC |  |
| --- | --- |
| Data collection |  |
| Space group | P 1 21 1 |
| Cell dimensions |  |
| a, b, c (Å) | 81.57, 113.23, 82.87 |
| $\alpha, \beta, \gamma$ (°) | 90.0, 108.80, 90.0 |
| Resolution (Å) | 66.84-2.3 (2.34-2.3) |
| Rsym or Rmerge | 0.149 (1.049) |
| I / $\sigma$(I) | 8.4 (0.6) |
| Completeness (%) | 99.8 (95.2) |
| Redundancy | 7 (6.4) |
| CC half | 0.991 (0.623) |
| Refinement |  |
| Resolution (Å) | 66.84-2.3 |
| No. reflections | 63174 |
| Rwork / Rfree | 0.22/0.27 |
| No. atoms |  |
| Protein | 9736 |
| Ligand/ion | 72 |
| Water | 82 |
| B-factors |  |
| Protein (min-max) | 49.78 (42.26 for L; 59.28 for C) |
| Ligand (min-max) | 55.95 (47.66 for H; 64.23 for A) |
| Water | 40.4 |
| R.m.s deviations |  |
| Bond lengths (Å) | 0.009 |
| Bond angles (°) | 1.28 |

Each dataset was collected from a single crystal. \*Values in parentheses are for highest-resolution shell.

